# Systemic hypoxia suppresses solid tumor growth

**DOI:** 10.64898/2026.02.09.704975

**Authors:** Ayush D. Midha, Brandon T. L. Chew, Benedict M. H. Choi, Jung Min Suh, Chris Carpenter, Alan H. Baik, Tej A. Joshi, Skyler Y. Blume, Augustinus G. Haribowo, Pedro Ruivo, Will R. Flanigan, Ankur Garg, Daniel D. Zhang, Vishvak Subramanyam, Rebecca Shuere, Youngho Seo, Henry VanBrocklin, Hani Goodarzi, Isha H. Jain

## Abstract

Local hypoxia is a hallmark of solid tumors and a negative prognostic factor in the progression and treatment of cancer. Here, we showed that systemic hypoxia, in contrast to localized tumor hypoxia, decreases tumor growth *in vivo* across multiple cancer types and preclinical models. The reduced tumor growth in systemic hypoxia was not explained by hypoglycemia, hypoinsulinemia, or HIF activation. Instead, metabolite profiling in tumors and tumor interstitial fluid revealed extensive perturbations in purine-related metabolites. Stable isotope tracing demonstrated that systemic hypoxia caused tumors to suppress *de novo* purine synthesis. Furthermore, tumors did not develop resistance to systemic hypoxia therapy, and when used in combination with chemotherapy or immunotherapy, systemic hypoxia dramatically suppressed tumor growth. Finally, we showed that systemic hypoxia can be achieved pharmacologically with the small molecule HypoxyStat. These findings challenge the long-held paradigm of hypoxia as a negative prognostic factor in cancer progression, and they suggest a potential therapeutic role for systemic hypoxia in suppressing solid tumor growth.

## INTRODUCTION

The unchecked growth and proliferation of cancer cells imposes substantial metabolic demands. Building new nucleotides, amino acids, and lipids force cancer cells to rely heavily on exogenous nutrients. Over a century ago, Otto Warburg demonstrated that cancer cells consume dramatically more glucose than their surrounding tissue^1^. Since this observation, it has become clear that depriving cancer cells of essential nutrients can suppress tumor growth. Indeed, interventions that starve tumors of key nutrients such as glucose^2–5^, arginine^6^, and methionine^7^ slow tumor growth in pre-clinical models. These findings support the paradigm that the availability of nutrients in the host environment shapes tumor metabolism and growth.

Among the nutrients in the host environment, oxygen is important for cancer cell proliferation. As the terminal electron acceptor in respiration, oxygen is required for mitochondrial ATP production. Moreover, respiration supports the synthesis of necessary anabolic substrates like serine^8^, aspartate^9–11^, and lipids^12^. Oxygen is also required for the production of unsaturated fatty acids, which maintain membrane homeostasis and organelle integrity^13,14^.

Nevertheless, local tumor hypoxia has been shown to facilitate tumor progression. In hypoxic cancer cells, activation of hypoxia-inducible factors (HIFs) increases glucose uptake^15^ and vascularization^16^. Local hypoxia also induces epigenetic reprogramming^17,18^ and confers resistance to radiotherapy^19–21^ and certain chemotherapies^22^. Indeed, tumor hypoxia is a negative prognostic factor for patient outcomes^23^.

Most studies on tumor hypoxia focus on *local* hypoxia caused by poor perfusion and rapid tumor expansion. However, the effects of *systemic* hypoxia remain unexplored. Systemic hypoxia, like that induced by altitude, initiates coordinated physiological responses including heightened erythropoiesis^24^, increased renal bicarbonate elimination^25^, and organ-specific metabolic rewiring^26^. These adaptations alter nutrient availability, hormone levels, and inter-organ communication^27^.

The effects of these global physiological changes on cancer growth and metabolism remain unknown. Here, we demonstrate that systemic hypoxia paradoxically suppresses cancer growth across multiple models and suppresses the synthesis of new purine nucleotides. These findings suggest that manipulating host nutrients—even those as fundamental as oxygen—can have anti-tumor effects, in contrast to the paradigm that hypoxia uniformly supports aggressive cancer progression.

## RESULTS

### Systemic hypoxia suppresses tumor growth across syngeneic and genetic models

To investigate the impact of systemic hypoxia on tumor proliferation, we used Panc02 cells, a pancreatic ductal adenocarcinoma (PDAC) cell line, to establish subcutaneous tumors in C57BL/6J mice. Mice were subsequently exposed to continuous atmospheric oxygen concentrations of 21% (normoxia), 11% (moderate hypoxia), or 8% (hypoxia) (**Fig. 1a**). We observed a significant reduction in tumor growth in mice subjected to systemic hypoxia (**Fig. 1b**), and tumors excised from hypoxic mice were visibly smaller compared to those from normoxic controls (**Fig. S1a**). In addition to subcutaneous models, systemic hypoxia also decreased tumor growth in orthotopic implants of E0771 cells, a syngeneic murine breast cancer line, in the mammary fat pads (**Fig. 1c**).

**Figure 1.**
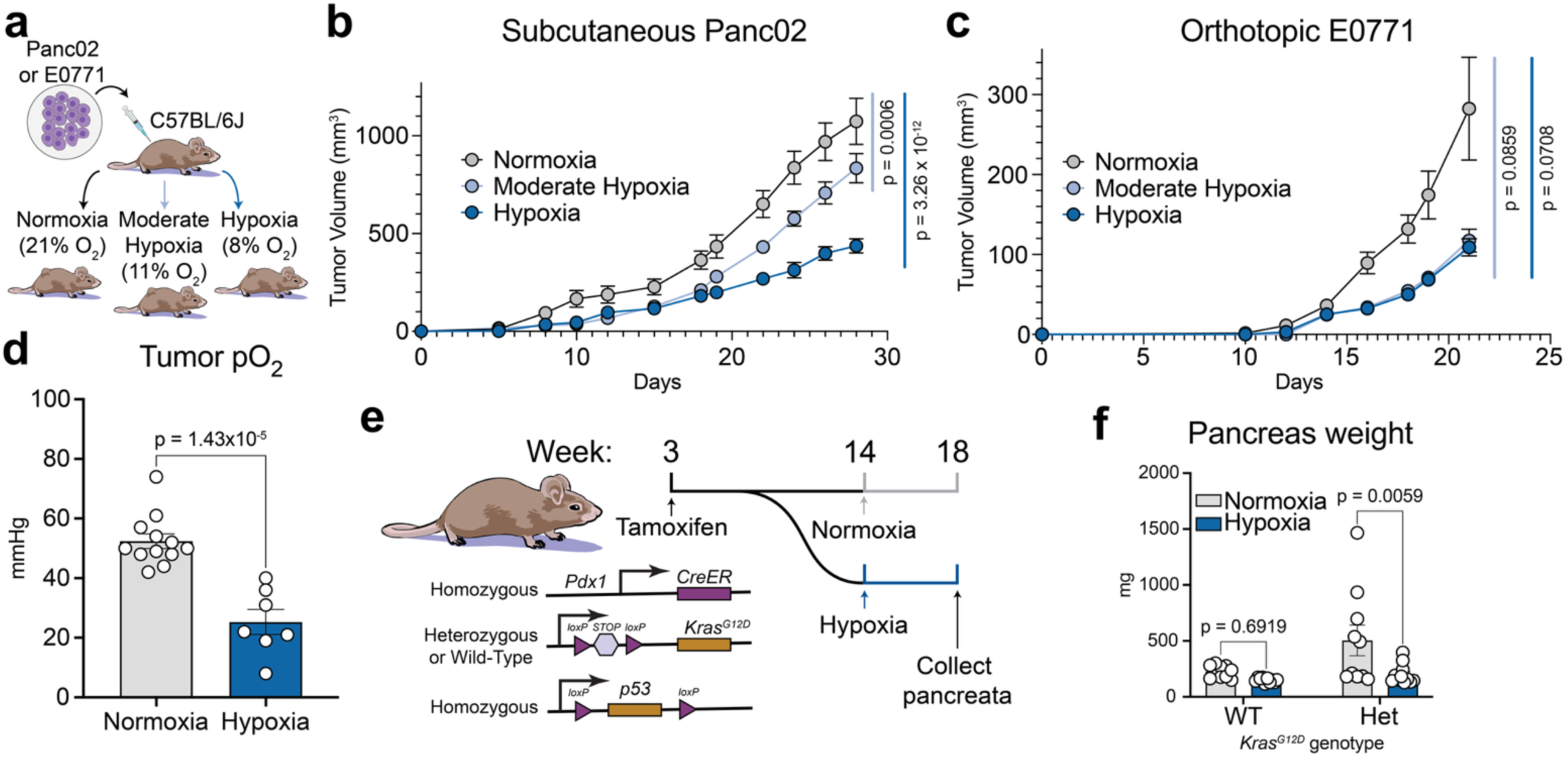
Systemic hypoxia suppresses tumor growth across syngeneic and genetic models. **a,** Schematic for subcutaneous syngeneic tumor models. **b,** Tumor growth of a subcutaneous implant of Panc02 cells in mice housed in normoxia (21% O_2_), moderate hypoxia (11% O_2_), or hypoxia (8% O_2_). Normoxia and hypoxia *n* = 7 mice, moderate hypoxia *n* = 6 mice. **c,** Tumor growth of an orthotopic implant of E0771 cells in mice housed in normoxia, moderate hypoxia, or hypoxia. *n* = 8 mice per group. **d,** Partial pressure of oxygen in tumors from mice in normoxia or hypoxia. Normoxia *n* = 12 mice, hypoxia *n* = 7 mice. **e,** Schematic for KPC tumor model. **f,** Pancreas weights from wild-type or heterozygous KPC mice housed in normoxia or hypoxia. Normoxia WT/Het and hypoxia WT *n* = 10 mice, hypoxia Het *n* = 11 mice. Data are presented as mean ± SEM. Comparisons were made using two-way ANOVA followed by Tukey’s multiple comparisons test **(b-c)**, two-tailed Student’s t-test **(d)**, or two-way ANOVA followed by Šídák’s multiple comparisons test **(f)**.

To ascertain the effect of systemic hypoxia on local tumor oxygenation, we directly measured the partial pressure of oxygen (pO_2_) within the tumors. We anesthetized and intubated mice to maintain their inhaled oxygen level, exposed their subcutaneous tumors, and used a Clark-type electrode to directly measure the pO_2_. As expected, tumors from mice housed in hypoxic conditions exhibited lower pO_2_ than those from normoxic counterparts (**Fig. 1d**).

We further extended our investigation to the *Kras^G12D^*;*p53^fl/fl^*;*Pdx1-CreER* (KPC) mouse model, a tamoxifen-inducible genetically engineered model of pancreatic cancer^28^. Ten weeks post-tamoxifen induction, mice that were wild-type (WT) or heterozygous (Het) for the Kras^G12D^ mutation were exposed to either hypoxic or normoxic conditions for four weeks. Pancreases were then collected, weighed, and analyzed by histopathology (**Fig. 1e**). Histopathological analysis revealed no pancreatic lesions or cancer development in any of the WT mice. In contrast, 75% of the Het mice developed pancreatic lesions (**Fig. S1b, Supp. Table 1**). The pancreas weight, a proxy for primary tumor growth, was significantly greater in normoxic Het mice than in hypoxic Het mice (**Fig. 1f**). Altogether, the above investigations suggested that systemic hypoxia suppresses solid tumor growth across multiple preclinical models.

### GENEVA reveals consistent but heterogeneous growth suppression across a diverse pool of cell line-derived xenografts

To investigate whether tumor suppression in systemic hypoxia is conserved across cancer types or shaped by lineage- or genotype-specific factors, we turned to our GENEVA (<u>gen</u>etically diverse and <u>e</u>ndogenously controlled phenotypic <u>v</u>ariation <u>a</u>ssay) platform^29^. GENEVA enables highly multiplexed xenograft experiments in which pooled human cancer cell lines form mosaic tumors, allowing lineage-resolved, single-cell profiling of both molecular and phenotypic responses *in vivo*. By assaying many genotypes simultaneously in the same host environment, GENEVA provides a scalable framework to detect both universal and context-specific effects of perturbations like hypoxia. The resulting *in vivo* phenotypic heterogeneity across cell lines, coupled with transcriptomic profiles at a single-cell resolution, allows us to dissect the molecular mechanisms underlying variable sensitivity to systemic hypoxia^29^.

To this end, we implanted immunocompromised NOD SCID gamma (NSG) mice with mosaic tumors composed of 20 human cancer cell lines spanning multiple tissue types and oncogenic backgrounds (**Fig. 2a**). Tumor-bearing mice were then exposed to either normoxic (21% O_2_) or hypoxic (8% O_2_) conditions for 8 days, after which tumors were resected and profiled by single-cell RNA-sequencing (Chromium Next GEM 3’ HT v3.1).

**Figure 2.**
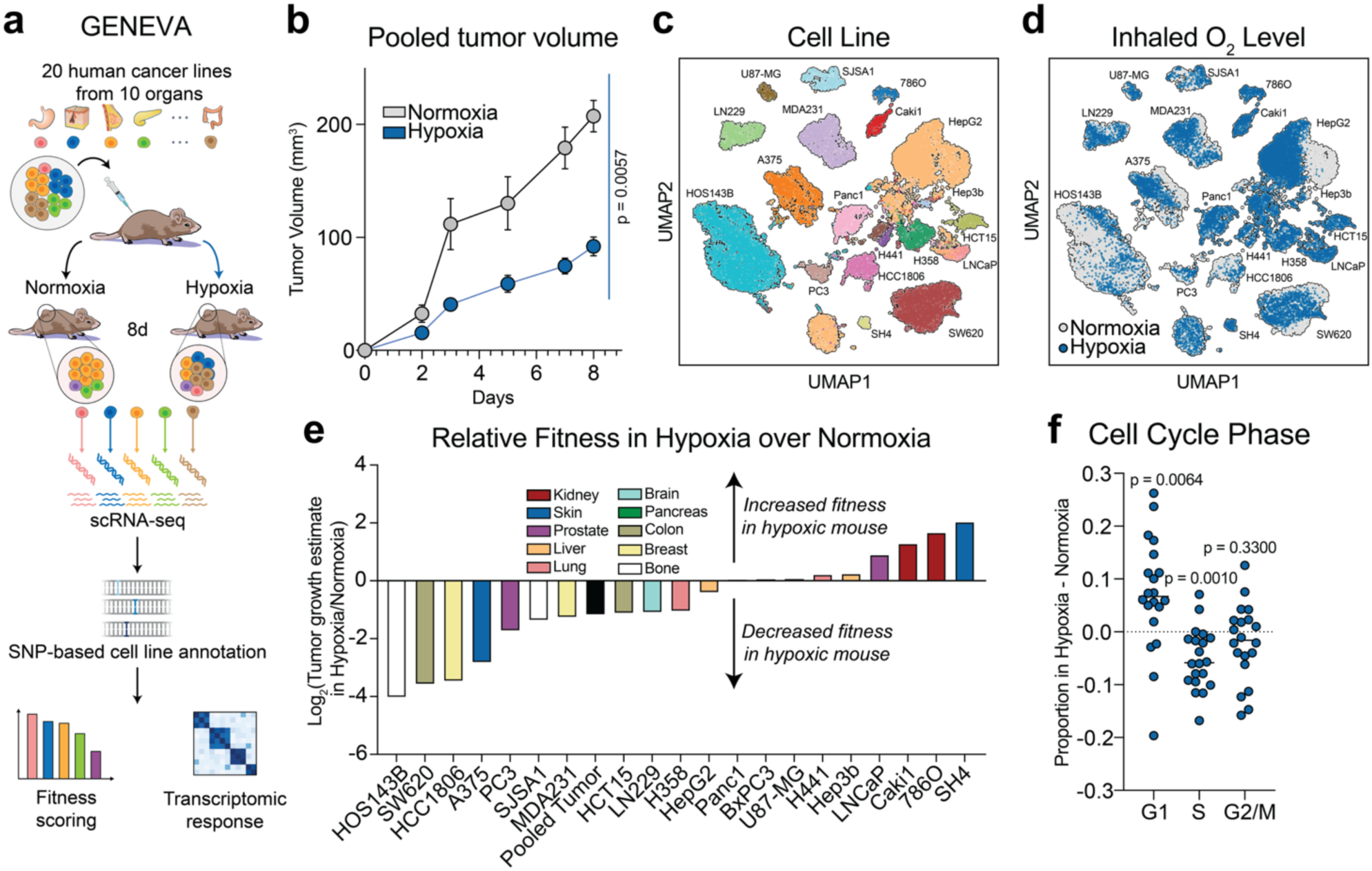
GENEVA reveals consistent but heterogeneous growth suppression across a diverse pool of cell line-derived xenografts. **a,** Schematic for the GENEVA experiment. **b,** GENEVA pooled tumor growth in mice housed in normoxia (21% O_2_) or hypoxia (8% O_2_). *n* = 4 mice per group. **c-d,** UMAP projection of single-cell transcriptomics from the GENEVA experiment with cells colored by cell line (c) or oxygen level (d). **e,** Relative fitness of each cell line from the GENEVA tumor in hypoxia compared to normoxia. *n* = 2 tumors per group. **f,** Difference in proportion of each cell line in G1, S, or G2/M phase between hypoxia and normoxia. *n* = 2 tumors per group. Data are presented as mean ± SEM. Comparisons were made using two-way ANOVA followed by Tukey’s multiple comparisons test **(b)** or Wilcoxon signed ranks test **(d)**.

Mirroring the effects observed in the syngeneic and GEMM models, systemic hypoxia led to a marked and significant decrease in mosaic tumor growth in the multiplexed xenograft setting (**Fig. 2b**). To estimate lineage-specific tumor growth, we leveraged SNP-based deconvolution to quantify the abundance of each cell line in the post-treatment tumors (**Fig. 2c-d**). We compared these proportions to each line’s initial representation in the injected pool and scaled the fold-change by total tumor volume to estimate cell line-specific tumor burden. Using this approach, we inferred a relative fitness score for each cell line under hypoxia versus normoxia (**Supp. Table 2**). While most lines exhibited reduced fitness under hypoxic conditions, a subset, including SH4 (melanoma), 786O, and Caki1 (renal cell carcinomas), showed increased relative growth (**Fig. 2e**), suggesting intrinsic resistance or metabolic adaptation to reduced oxygen availability.

To further explore this observation, we scored cell cycle states using hallmark gene signatures. Across most lines, hypoxia induced a shift toward G1 arrest and reduced S-phase occupancy (**Fig. 2f, S1c**), consistent with a general suppression of proliferation. Yet, the extent of this cell cycle shift varied by lineage and correlated with differential tumor growth. This highlights cell intrinsic molecular heterogeneity that leads to variation in response to systemic hypoxia.

### Systemic hypoxia suppresses tumor growth independently of hypoglycemia, hypoinsulinemia, or HIF activation

Next, we explored potential mechanisms by which systemic hypoxia could suppress tumor growth. Systemic hypoxia induces hypoglycemia^26,30^, and interventions that deprive tumors of glucose, such as brown fat activation^3–5^ and low-glycemic diets^2^, have been shown to suppress tumor growth. Consistent with our prior observations, hypoxia exposure lowered blood glucose of tumor-bearing mice (**Fig. 3a**).

**Figure 3.**
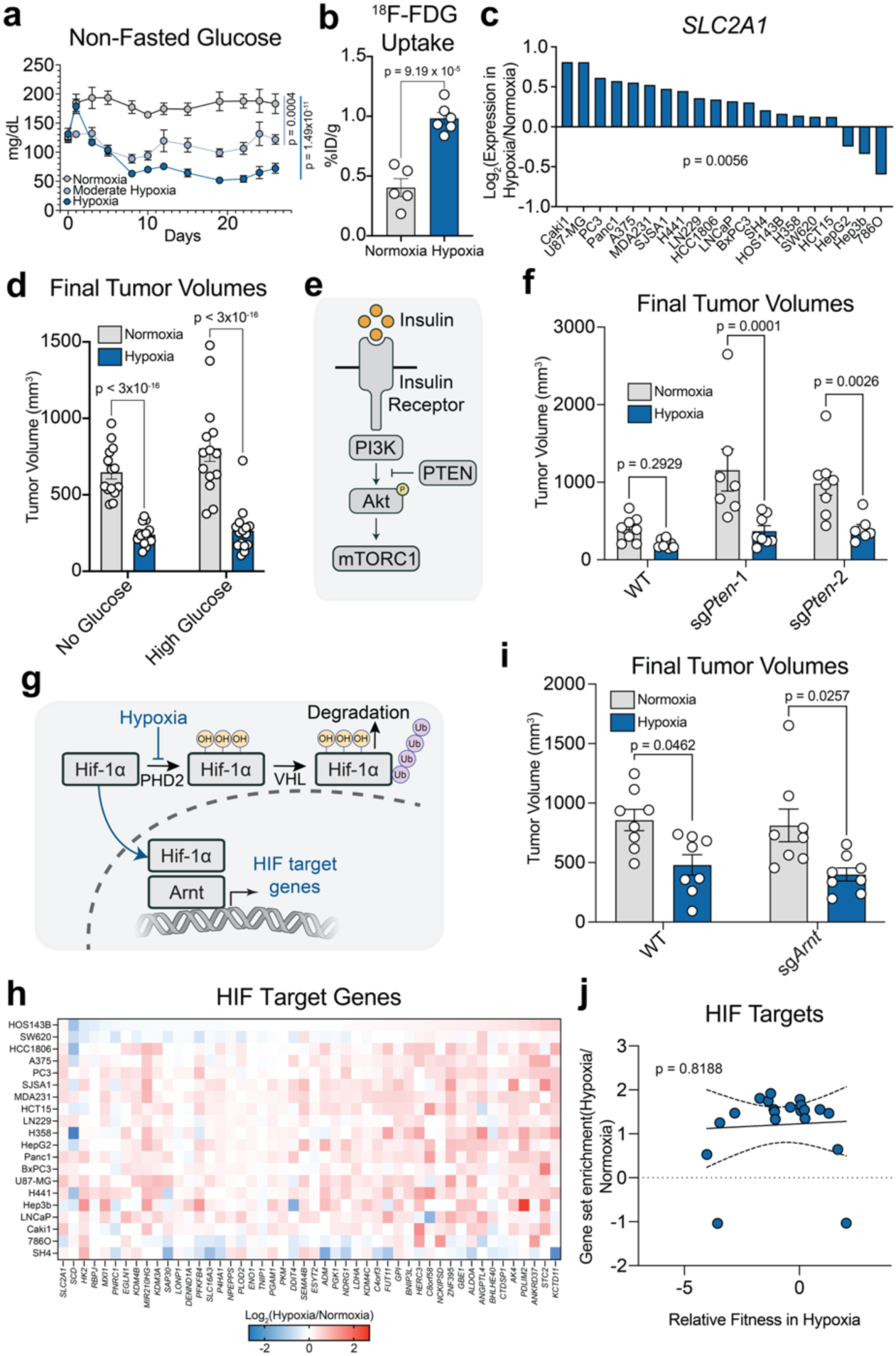
Systemic hypoxia suppresses tumor growth independently of hypoglycemia, hypoinsulinemia, or HIF activation. **a,** Non-fasted blood glucose levels of tumor-bearing mice housed in normoxia (21% O_2_), moderate hypoxia (11% O_2_), or hypoxia (8% O_2_). Normoxia and hypoxia *n* = 7 mice, moderate hypoxia *n* = 6 mice. **b,** ^18^F-FDG uptake by tumors, measured as the percent of the injected dose normalized to tumor weight (%ID/g), from mice housed in normoxia or hypoxia. Normoxia *n* = 5 mice, hypoxia *n* = 6 mice. **c,** Fold changes between hypoxia and normoxia in the expression of *SLC2A1* across the different cancer cell lines from GENEVA tumors. **d,** Final volumes of Panc02 tumors in mice housed in normoxia or hypoxia and supplied water with no glucose or 30% glucose. Hypoxia with 30% glucose *n* = 16 mice, every other group *n* = 14 mice. **e,** Schematic showing the regulation of insulin signaling pathway by PTEN. **f,** Final volumes of tumors composed of WT Panc02 cells and two *Pten*-knockout clones. Normoxia WT, normoxia sg*Pten*-2, hypoxia WT, and hypoxia sg*Pten*-1 *n* = 8 mice, normoxia sg*Pten*-1 and hypoxia sg*Pten*-2 *n* = 7 mice. **g,** Schematic of Hif-1⍺ pathway. **h,** Fold change in expression of HIF target genes in different cancer cell lines from GENEVA tumors that were implanted into mice housed in normoxia or hypoxia. **i,** Final volumes of tumors composed of WT Panc02 cells and an *Arnt*-knockout clone. *n* = 8 mice. **j,** Correlation between the change in GSEA normalized enrichment scores for HIF target genes between hypoxia and normoxia and relative fitness in hypoxia. Data are presented as mean ± SEM. Comparisons were made using two-way ANOVA followed by Tukey’s multiple comparisons test **(a, d, f, i)**, two-tailed Student’s t-test **(b)**, or Wilcoxon signed ranks test **(c)**. A linear regression was conducted to test the significance of the relationship between fold change in gene expression and relative fitness in hypoxia **(j)**.

To assess the impact of hypoxia on tumor glucose uptake, we injected mice with the radioactive glucose analogue ^18^F-fluorodeoxyglucose (FDG). After housing the mice in their respective O_2_ levels for 1 hour, we dissected tumors and quantified the gamma radiation in each tumor. Intriguingly, tumors in hypoxic mice took up twice as much ^18^F-FDG as those in normoxic mice (**Fig. 3b**). Furthermore, in the GENEVA tumors, hypoxia upregulated the expression of *SLC2A1*, which encodes the glucose transporter GLUT1, the major driver of FDG uptake in most cell lines (**Fig. 3c**). Across cell lines, there was no relationship between changes in *SLC2A1* expression and fitness in hypoxia (**Fig. S2a**). Together, these data suggest that while systemic hypoxia induces hypoglycemia, most cancer cells compensate by increasing their uptake of glucose. To confirm that tumor suppression in hypoxic mice was independent of hypoglycemia, we gave mice with water supplemented with 30% glucose. Glucose supplementation slightly increased tumor growth in normoxic mice but had no discernible effect in hypoxic mice (**Fig. 3d, S2b**).

Systemic hypoxia also lowers circulating insulin levels^26^, so we also considered whether decreased insulin signaling could account for tumor suppression under hypoxic conditions. The binding of insulin to the insulin receptor on cancer cells activates PI3K, initiating a pro-growth signaling cascade^31^. PI3K is negatively regulated by PTEN, so PTEN-deficient cells exhibit constitutive insulin signaling (**Fig. 3e**). Cancer cells with constitutive insulin signaling are known to be resistant to the tumor-suppressive effects of dietary restriction^32^. To determine whether constitutive insulin signaling could override the effects of systemic hypoxia, we generated single-clone knockouts of *Pten* in Panc02 cells (**Fig. S2c**). In normoxic mice, *Pten*-deficient tumors grew more rapidly than wild-type tumors. However, hypoxia treatment still significantly decreased the growth of *Pten*-deficient tumors, suggesting that hypoxia’s suppressive effect was independent of insulin signaling (**Fig. 3f**, **Fig. S2d**).

Another potential mechanism for tumor suppression in hypoxia is the activation of hypoxia-inducible factor (HIF). In normoxic cells, HIF-1⍺ is hydroxylated by prolyl hydroxylase domain (PHD) enzymes in an oxygen-dependent reaction. This hydroxylation marks HIF-1⍺ for VHL-mediated ubiquitination, leading to its degradation. In hypoxic conditions, HIF-1⍺ is not degraded; it translocates to the nucleus, complexes with HIF-1β, and drives the transcription of genes involved in adaptation to hypoxia (**Fig. 3g**)^33–36^. HIF-inducible transcripts can have context-dependent effects on tumor growth. While some HIF transcripts promote glycolysis and angiogenesis, potentially fostering tumor growth, HIF activation can also induce apoptosis^37^. For instance, in a hepatocellular carcinoma model, HIF inhibition suppresses tumor growth^38^, but in an AML model, HIF activation has been shown to suppress tumor growth^39^.

In our GENEVA experiment, hypoxia treatment increased the expression of most consensus HIF target genes^40^ in the majority of cell lines (**Fig. 3h**). To determine whether activation of HIF target genes is necessary for the observed tumor growth suppression, we generated single-clone knockouts of *Arnt*, the gene encoding HIF-1β (**Fig. S2e-f**). In both normoxia and hypoxia, there was no significant difference in tumor growth between wild-type and *Arnt*-deficient cells, indicating that HIF-dependent transcription in cancer cells is not necessary for tumor suppression in hypoxia (**Fig. 3i**, **Fig. S2g**). In the GENEVA model, there was no significant association between the activation of HIF targets and relative fitness in hypoxia (**Fig. 3j**). Altogether, these findings confirm that the tumor suppressive effects of systemic hypoxia are not dependent on tumor-intrinsic HIF stabilization.

### Systemic hypoxia reprograms tumor nucleotide metabolism

After ruling out these potential mechanisms of tumor suppression, we took a more unbiased approach to investigate the metabolic basis of slowed tumor growth in systemic hypoxia. To this end, we profiled the metabolites in tumor homogenates and tumor interstitial fluid (TIF) of subcutaneous Panc02 tumors from mice housed in normoxia and hypoxia. Nucleotide intermediates and nucleotide derivatives were among the most significantly depleted metabolites in tumor homogenates from mice exposed to hypoxia (**Fig. 4a**). TIF from hypoxic tumors also exhibited lower nucleotide levels and was enriched with the nucleotide breakdown product hypoxanthine (**Fig. 4b**). Tumors are known to exhibit substantially higher nucleotide levels than other cells^41^, and depletion or imbalance of nucleotides can impair DNA replication^42,43^.

**Figure 4.**
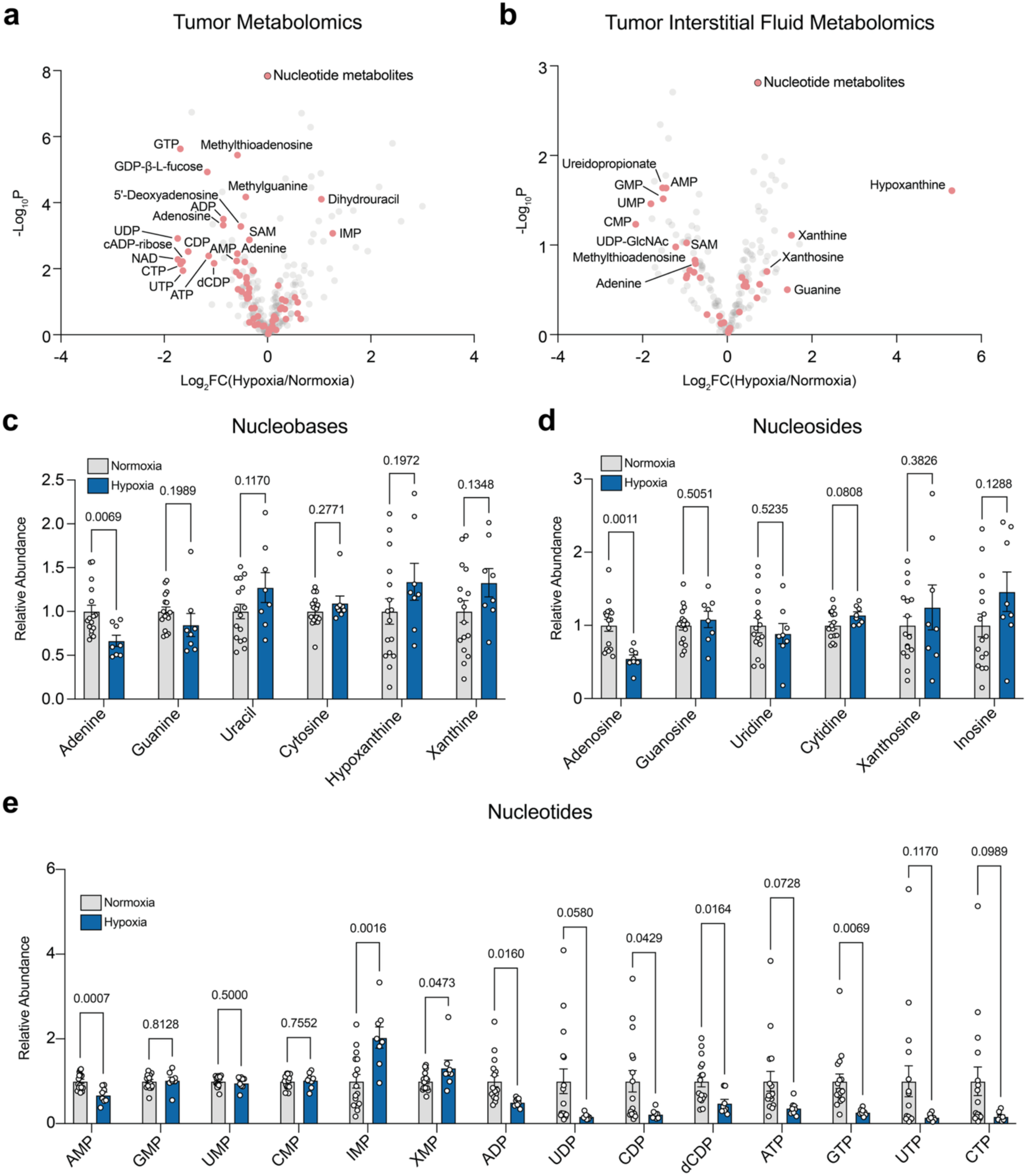
Systemic hypoxia reprograms tumor nucleotide metabolism. **a,** Changes in abundance of metabolites in tumor homogenates from mice housed in normoxia (21% O_2_) or hypoxia (8% O_2_). Normoxia *n* = 16 mice, hypoxia *n* = 8 mice. **b,** Changes in abundance of metabolites in tumor interstitial fluid from mice housed in normoxia or hypoxia. Normoxia *n* = 7 mice, hypoxia *n* = 6 mice. **c-e,** Fold changes in nucleobases **(c)**, nucleosides **(d)**, and nucleotides **(e)** in tumor homogenates between mice housed in normoxia or hypoxia. Data are presented as mean ± SEM. Comparisons were made using two-tailed Student’s t-tests.

From the tumor metabolomics, we analyzed levels of nucleobases, nucleosides (nucleobase + ribose sugar), and nucleotides (nucleoside + phosphates). Nucleobase and nucleoside levels did not vary significantly between hypoxic and normoxic tumors, with the notable exception of adenine and adenosine (**Fig. 4c-d**). Among mononucleotides, adenosine monophosphate (AMP) was depleted in hypoxic tumors, while the purine intermediate inosine monophosphate (IMP) was enriched in hypoxic tumors. Nearly all measured dinucleotides and trinucleotides were depleted in hypoxic tumors (**Fig. 4e**). Altogether, hypoxia induced the depletion of purine and pyrimidine synthesis products (**Fig. S3a-b**) and the accumulation of some salvage/breakdown intermediates (**Fig. 4c-d**). The specific depletion of adenine, adenosine, and AMP among nucleobases, nucleosides, and mononucleotides in hypoxia suggested a unique deficit in the synthesis of purines.

### Cancer cells susceptible to hypoxia therapy suppress de novo purine synthesis

Purine nucleotides can be assembled through either the *de novo* synthesis pathway or through the salvage of nucleobase precursors. *De novo* synthesis is energetically demanding and requires multiple enzymes to incorporate components derived from aspartate, glycine, glutamine, and one-carbon donors into the purine ring^44^. Salvage, in contrast, involves the uptake of nucleobases and their conjugation with phosphoribosyl pyrophosphate (PRPP) from the pentose phosphate pathway (**Fig. 5a**). Previous work has shown that in hypoxic cells in culture, aspartate can become limiting, diminishing *de novo* purine synthesis^10,45^. However, in the hypoxic tumor homogenates, aspartate levels were not depleted, suggesting that aspartate deprivation does not explain the depletion of purine nucleotides (**Fig. S4a**).

**Figure 5.**
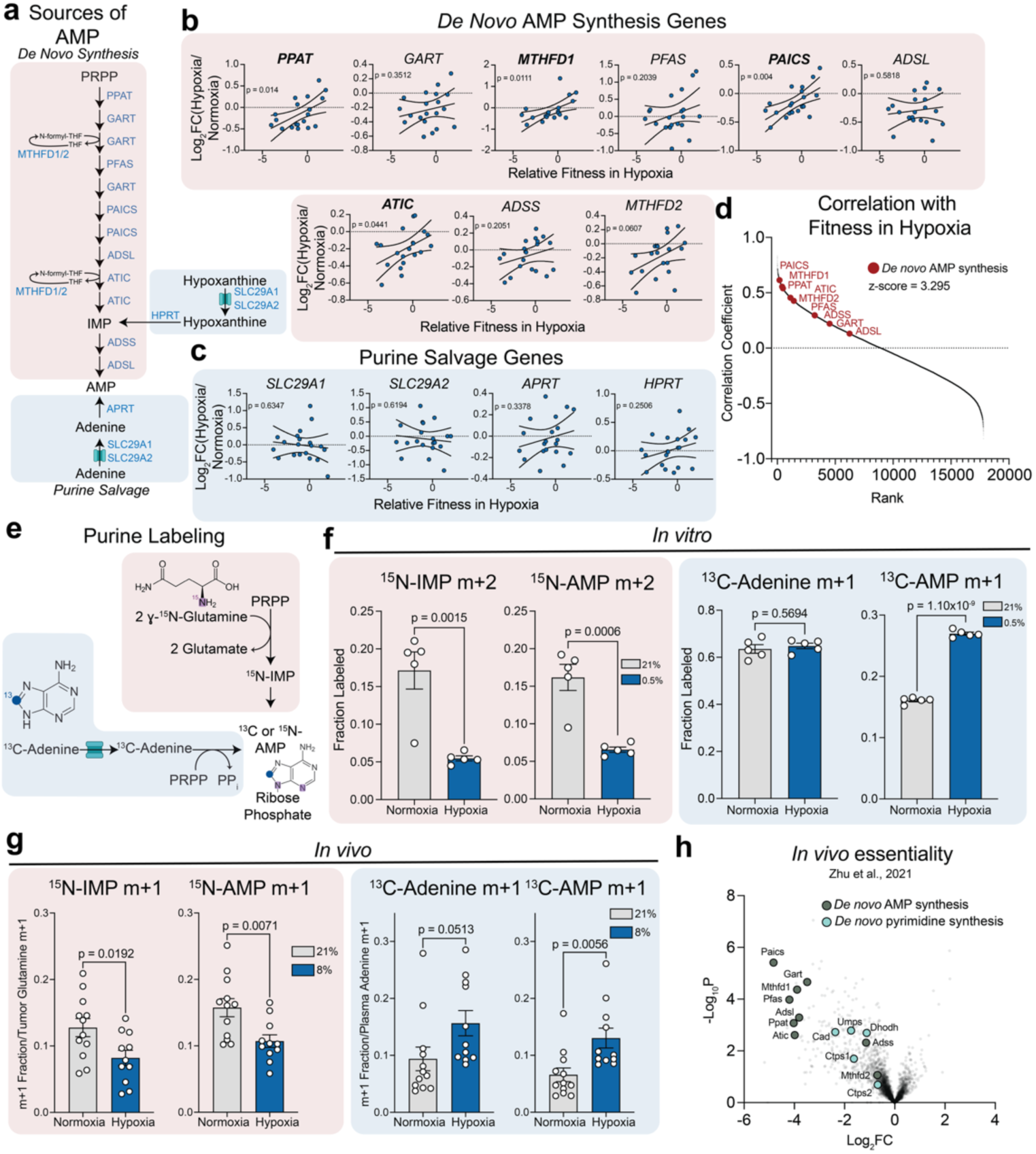
Cancer cells susceptible to hypoxia therapy suppress *de novo* purine synthesis. **a,** Schematic of *de novo* AMP synthesis and purine salvage. **b-c,** Correlation between fold change in gene expression between hypoxia (8% O_2_) and normoxia (21% O_2_) and the relative fitness in hypoxia of each cell line from GENEVA tumors. Selected genes belong to the *de novo* AMP synthesis (**b**) or purine salvage (**c**) pathways. *n* = 2 tumors in each O_2_, *n* = 20 cell lines. **d,** Ranked correlation coefficients between fold change in gene expression and relative fitness in hypoxia. *De novo* AMP synthesis genes are labeled in red. **e,** Schematic of dual ɣ-^15^N-glutamine/8-^13^C-adenine labeling experiment. **f,** Fractional labeling of ^15^N-IMP, ^15^N-AMP, ^13^C-adenine, and ^13^C-AMP in Panc02 cells after 6 hours of incubation with ɣ-^15^N-glutamine/8-^13^C-adenine media in normoxia (21% O_2_) or hypoxia (0.5% O_2_). *n* = 5. **g,** Fractional labeling of intra-tumor ^15^N-IMP and ^15^N-AMP normalized to intra-tumor ^15^N-glutamine fractional labeling. Fractional labeling of intra-tumor ^13^C-adenine and ^13^C-AMP normalized to plasma ^13^C-adenine fractional labeling. Tumors were composed of Panc02 cells and were implanted into mice housed in normoxia (21% O_2_) or hypoxia (8% O_2_). Mice were infused with a ɣ-^15^N-glutamine/8-^13^C-adenine solution for 5 hours in the given O_2_ tension. Normoxia *n* = 12 mice, hypoxia *n* = 11 mice. **h,** Essentiality scores of a library of metabolism-focused genes in subcutaneous pancreatic cancers implanted into C57BL/6J mice. Adapted from *Zhu et al., 2021*^48^. Data are presented as mean ± SEM. Linear regressions were conducted to test the significance of the relationship between fold change in gene expression and relative fitness in hypoxia **(b-c)**, and correlation coefficients were calculated using Pearson correlation **(d)**. The z-score for enrichment of *de novo* AMP synthesis genes was calculated using iPAGE **(d)**. Comparisons were made using two-tailed Student’s t-tests **(f-g)**.

Next, we investigated the relationship between purine synthesis gene expression and *in vivo* cancer cell fitness in systemic hypoxia. Because GENEVA provides high-resolution molecular profiles across a genetically diverse pool of cancer models, it offers a unique opportunity to leverage natural phenotypic heterogeneity to uncover molecular determinants of response. Using single-cell transcriptomics from GENEVA mosaic tumors, we measured the correlation between hypoxia-induced transcriptional changes in purine metabolism genes and lineage-specific fitness in hypoxia. Changes in expression of several key *de novo* purine synthesis genes, i.e., *PPAT, MTHFD1, PAICS, and ATIC*, were significantly and positively associated with relative fitness in hypoxia (**Fig. 5b**). Most cell lines which showed slower growth in hypoxic mice also downregulated these genes, whereas hypoxia-tolerant lines maintained expression of these biosynthetic enzymes (**Fig. S4b-c**). In contrast, expression of purine salvage pathway genes showed no relationship with fitness (**Fig. 5c**), and expression of *de novo* pyrimidine synthesis genes were not consistently altered (**Fig. S4b**). Across the genome, *de novo* purine synthesis genes were significantly enriched among genes whose expression was positively associated with fitness in hypoxia (**Fig. 5d, Fig. S4d**).

Based on these findings, we proceeded to directly measure *de novo* purine synthesis and purine salvage. To quantify the relative contributions of these pathways, we cultured cells with both ɣ-^15^N−glutamine and 8−^13^C-adenine (**Fig. 5e**). Using this approach, *de novo* synthesized purines would carry a ^15^N label and salvaged purines would carry a ^13^C label. In Panc02 cells *in vitro*, hypoxia decreased the fraction of IMP and AMP with ^15^N labels and increased the fraction of AMP carrying a ^13^C label, indicating a suppression of *de novo* purine synthesis and an increased reliance on salvage (**Fig. 5f**).

We performed a similar experiment *in vivo*, continuously infusing a solution of ɣ-^15^N−glutamine and 8−^13^C-adenine into mice with subcutaneous Panc02 tumors. The infusion labeled a substantial fraction of circulating glutamine and adenine pools, as well as the intratumor glutamine pool (**Fig. S4c**). To quantify the contribution of *de novo* purine synthesis, we measured the fractional labeling of intratumor ^15^N-IMP and ^15^N-AMP normalized to intratumor ^15^N-glutamine labeling. Normalized ^15^N labeling of IMP and AMP was lower in hypoxic tumors (**Fig. 5g**, **Fig. S4d-e**), and ^15^N labeling of UMP and GMP was not altered in hypoxia (**Fig. S4f-i**). To quantify the contribution of adenine salvage, we measured the fractional labeling of intratumor ^13^C-adenine and ^13^C-AMP, normalized to circulating ^13^C-adenine labeling. Normalized ^13^C labeling of AMP was significantly higher in hypoxic tumors (**Fig. 5g**). Altogether, both the *in vitro* and *in vivo* labeling experiments indicate that hypoxia suppressed *de novo* purine synthesis and increased the contribution of purine salvage to intracellular AMP pools. These findings are consistent with prior work showing increased dependence on purine salvage in states of bioenergetic stress^46,47^.

Of note, impaired *de novo* purine synthesis is sufficient to suppress tumor growth. A previously published CRISPR-knockout screen of metabolic genes in subcutaneous pancreatic tumors revealed that genes involved in *de novo* purine synthesis were some of the most essential genes for tumor growth *in vivo*^48^ (**Fig. 5h**). These genes were significantly Less essential *in vitro*, suggesting that the *in vivo* tumor environment uniquely renders proliferating cells dependent on *de novo* synthesis of purine nucleotides. Altogether, these results suggest that systemic hypoxia diminishes *de novo* purine synthesis in tumors, contributing to the suppression of tumor growth.

Next, we investigated potential mechanisms suppressing *de novo* purine synthesis in hypoxic tumors. Across all cell lines in the GENEVA tumor, changes in the expression of the *de novo* purine synthesis genes *PPAT*, *MTHFD1*, *MTHFD2*, *PAICS*, and *ATIC* in hypoxia were correlated with changes in the expression of most Myc targets^49^ (**Fig. S5a**). Myc is a known transcriptional driver of several purine synthesis genes^50–52^. Moreover, transcriptional changes in Myc targets were correlated with fitness in hypoxia, indicating that the cell lines that grew the least in hypoxia exhibited increased suppression of Myc activity (**Fig. S5b**). Further investigation is required to dissect the effects of systemic hypoxia on Myc activity and the role of Myc in regulating *de novo* purine synthesis.

### Systemic hypoxia is a potent anti-cancer therapy that synergizes with established first-line cancer therapies

Clinically, tumor-acquired resistance to therapy often compromises the efficacy of long-term treatments. Thus, we sought to determine whether tumors develop resistance to systemic hypoxia therapy by serially re-implanting tumors *in vivo* from mice housed in either normoxia or hypoxia. Previous work using this technique has demonstrated that tumors can adapt after several passages^53,54^. We implanted Panc02 cells into the flank of male C57Bl/6J mice and housed the mice in either 21% or 8% O_2_. Tumors were collected after approximately one month of growth, dissociated into single cells, expanded *in vitro*, and reimplanted into new mice. This process was repeated four times (**Fig. 6a**). Over repeated passages, tumors in hypoxic mice consistently grew slower than tumors in normoxic mice, suggesting that no overt resistance phenotype emerged (**Fig. 6b**).

**Figure 6.**
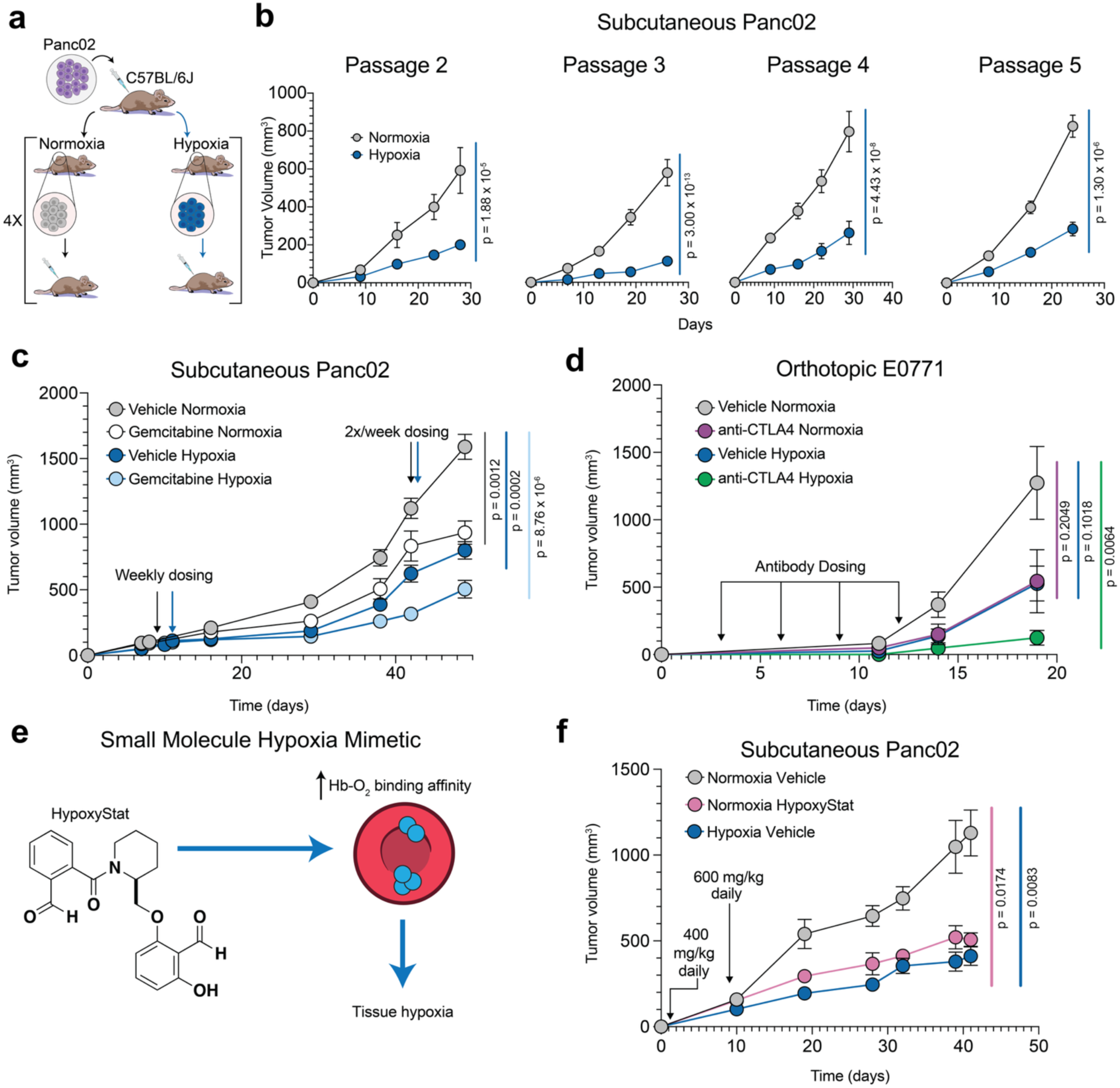
Systemic hypoxia is a potent anti-cancer therapy that synergizes with established first-line cancer therapies. **a,** Schematic of tumor reimplantation experiment. **b,** Growth of subcutaneous tumors of Panc02 cells that have undergone 2-5 passages *in vivo* in mice housed in normoxia (21% O_2_) or hypoxia (8% O_2_). Passage 2 *n* = 9 mice, Passage 3 normoxia *n* = 8 mice, Passage 3 hypoxia *n* = 7 mice, Passage 4 normoxia *n* = 8 mice, Passage 4 hypoxia *n* = 9 mice, Passage 5 *n* = 12 mice. **b,** Growth of subcutaneous tumors of Panc02 cells in mice that were housed in normoxia or hypoxia and treated with either vehicle or gemcitabine. *n* = 8 mice. **c,** Growth of orthotopic tumors of E0771 cells in mice that were housed in normoxia or hypoxia and treated with either vehicle or anti-CTLA4 antibody. Vehicle normoxia *n* = 12 mice, anti-CTLA4 normoxia *n* = 13 mice, Vehicle hypoxia *n* = 10 mice, anti-CTLA4 hypoxia *n* = 10 mice. **d,** Schematic of manipulation of hemoglobin (Hb)-O_2_ binding by HypoxyStat. **e,** Growth of subcutaneous Panc02 tumors in mice that were housed in normoxia or hypoxia and treated with either a vehicle or HypoxyStat. Normoxia vehicle *n* = 5 mice, normoxia HypoxyStat *n* = 7 mice, hypoxia Vehicle *n* = 8 mice. Data are presented as mean ± SEM. Comparisons were made using two-way ANOVA followed by either Šídák’s multiple comparisons test **(a)** or Tukey’s multiple comparisons test **(b, c, e)**.

To demonstrate the therapeutic potential of systemic hypoxia, we deployed systemic hypoxia as an adjuvant therapy alongside clinically relevant treatments. We first combined hypoxia treatment with gemcitabine monotherapy, a common first-line treatment for metastatic PDAC until 2011^55^. We subcutaneously implanted Panc02 cells in the flanks of male C57BL/6J mice and exposed the mice to either hypoxia or normoxia. Once tumors reached 100 mm^3^ in size, mice received 100 mg/kg gemcitabine or saline via intraperitoneal injection once weekly. After three weeks, the injections were escalated to twice weekly at the same dose. Gemcitabine and hypoxia treatment alone decreased tumor growth, and the combination of the two treatments further slowed tumor growth (**Fig. 6c**).

Next, we combined systemic hypoxia with an immune checkpoint inhibitor (ICI). We orthotopically injected E0771 cells into the mammary fat pads of female C57BL/6J mice and exposed the mice to either hypoxia or normoxia. Mice were injected with 100 µg of either an anti-mouse Ctla4 antibody or an IgG-2b isotype control antibody on days 3, 6, 9, and 12 after tumor implantation. Systemic hypoxia plus isotype control decreased tumor growth to a similar extent as ICI in normoxia alone, and systemic hypoxia plus ICI further decreased tumor growth, nearly completely abolishing cancer progression (**Fig. 6d**). Together, these results suggest that hypoxia synergizes with clinically relevant treatment modalities.

Inhaled hypoxia presents challenges for clinical implementation: patients would either need to be confined to hypoxic rooms or carry hypoxic gas mixtures delivered by nasal cannula. Both approaches have notable drawbacks, including decreased quality of life and a significant risk of equipment malfunction. To circumvent these issues and demonstrate the clinical translatability of hypoxia therapy, we used HypoxyStat^56^, our small molecule hypoxia-mimetic that decreases oxygen off-loading from hemoglobin, thereby serving as a small molecule form of hypoxia therapy. We treated male C57BL/6J mice carrying subcutaneous Panc02 tumors with either 400 mg/kg of HypoxyStat or a vehicle control daily (**Fig. 6e**). After 1 week, the HypoxyStat dose was escalated to 600 mg/kg. HypoxyStat dosing was sufficient to decrease tumor growth relative to the vehicle control and to a comparable degree as inhaled hypoxia (**Fig. 6f**).

Given our findings, we asked whether living at high altitude may confer a lower risk of cancer mortality, as has been previously suggested in various contexts^57–60^. To measure the association between elevation and cancer mortality, we first generated congruent per-county elevation and age-adjusted pan-cancer mortality rate (AAMR) maps using the National Oceanic and Atmospheric Administration’s ETOPO 2022 Global Relief Model and the CDC’s cancer data and statistics, respectively (**Fig. S6a**). Higher altitude was significantly negatively correlated with AAMR (**Fig. S6b**). Even after performing a multivariate regression and adjusting for median income and the prevalence of obesity and diabetes, the negative correlation between elevation and AAMR remained statistically significant (**Fig. S6c**). This epidemiological data supplies orthogonal evidence that systemic hypoxia suppresses tumor progression in humans.

## DISCUSSION

Our findings show that inhaled hypoxia suppresses primary tumor growth. This tumor suppression persists in conditions where glucose availability is increased, insulin signaling is activated, and HIF activation is impaired. Using metabolomics, single-cell transcriptomics, and stable-isotope labeled tracer experiments, we found that hypoxia inhibits *de novo* purine synthesis. Furthermore, the tumor suppressive effects of hypoxia synergize with the chemotherapy gemcitabine and immune checkpoint blockade therapy. Altogether, our findings may explain the observation that higher altitude counties in the United States generally have lower cancer mortality.

The tumor suppressive effects of systemic hypoxia vary by cancer types. Most experiments were conducted using Panc02 cells, which harbor the oncogenic *Kras^G12D^* mutation. Kras activation in PDAC drives a signaling cascade that increases nucleotide pools to support proliferation^61^, and this phenotype was dramatically reversed by systemic hypoxia (**Fig. 4e**). Strikingly, we identified a few cell lines (SH4, 786O, and Caki1) that grew better in hypoxic mice than in normoxic mice. Interestingly, two of these cell lines (Caki1 and 786O) are derived from clear cell renal cell carcinomas (ccRCC). The vast majority of ccRCC tumors exhibit the loss of the tumor suppressor VHL, resulting in persistent HIF activation^62^. However, Caki1 cells express wild-type *VHL*, suggesting that resistance to hypoxia-induced tumor suppression in ccRCC tumors may emerge from other metabolic features independent of VHL loss. Whereas most cell lines downregulated *de novo* purine synthesis genes in hypoxia, the two ccRCC cell lines evaded this transcriptional response (**Fig. 5b, Fig. S4a**).

We found that hypoxia-induced suppression of *de novo* purine synthesis was associated with the downregulation of Myc targets. In multiple contexts, Myc binds to the promoter regions of *de novo* purine synthesis genes^50–52^. Suppression of *de novo* purine synthesis in hypoxia might be a protective adaptation because building new AMP molecules is energetically costly, requiring 4 molecules of ATP and 1 molecule of GTP. Indeed, nutrient deprivation is known to suppress Myc activity^63^, and this suppression is protective for survival^64^. Despite being adaptive for cell survival, suppression of Myc activity would still slow overall tumor progression. Further work is required to dissect the role of decreased Myc activity in mediating the tumor suppressive effects of systemic hypoxia.

Intriguingly, we found that hypoxia diminished *de novo* purine synthesis in PDAC cells both *in vitro* and *in vivo*. Previous work has demonstrated that *de novo* purine synthesis is more essential in PDAC tumors *in vivo* than in PDAC cells in culture^48^. This discrepancy may arise because tumors *in vivo* are poorly vascularized and deprived of exogenous substrates required to bypass *de novo* purine synthesis. The purine salvage substrate hypoxanthine is abundant in tumor interstitial fluid^65^, but this observation may reflect the intra-tumor turnover of nucleotide pools rather than the availability of new purine intermediates. Indeed, we found that the tumor interstitial fluid of hypoxic tumors contained greater hypoxanthine levels than that of normoxic tumors, but this was insufficient to overcome the suppression of *de novo* purine synthesis *in vivo* (**Fig. 4b**).

Our findings present a stark contrast to the observation that local tumor hypoxia promotes aggressive tumor growth. When a tumor contains hypoxic neighborhoods, the hypoxic cells promote angiogenesis^16^, nutrient uptake^15^, and resistance to cancer-directed therapies^19–22^. Meanwhile, well-oxygenated pockets can likely undertake energetically demanding processes like nucleotide synthesis while supplying excess salvageable substrates to the poorly oxygenated regions. However, our work underlines that when the entire tumor (or entire host) is hypoxic, there is tumor-wide depletion of nucleotides. This disrupted division of labor in systemic hypoxia may also manifest for other energetically demanding processes, like cholesterol synthesis and lipogenesis. Indeed, hypoxic cancer cells become reliant on salvaging exogenous sources of cholesterol^66^ and fatty acids^67^.

The effects of systemic hypoxia on tumor growth are likely multifactorial because systemic hypoxia induces global physiological and metabolic changes. Future work is required to identify the effects of systemic hypoxia on tumor-intrinsic and tumor-extrinsic growth signaling, the availability of exogenous nutrients, and immune surveillance of tumors. Dissecting these mechanisms will enable the development of precision cancer therapies that target different pathways to maximize tumor suppression.

Altogether, our findings highlight the potential therapeutic role of systemic hypoxia as an adjuvant in cancer therapy. Our work on hypoxia as a therapy has gained traction for mitochondrial disease^68,69^, cardio-metabolic disease^26,30,70^, and aging^71,72^. The fact that communities at altitude have thrived for generations with lower mortality from cardiometabolic disease and cancer supports the translational potential of these findings into human patients. Moreover, the development of small molecules that increase hemoglobin’s binding affinity to oxygen has brought closer the clinical application of hypoxia as a therapy^56,73^. Nonetheless, optimization of these compounds is required to expand the therapeutic window. In addition, hypoxia can introduce health risks^72^, including pulmonary hypertension^74^, erythrocytosis^75^, and cerebral and pulmonary edema^76^. Further investigation of these side effects is required to limit the risks while advancing the potential of hypoxia as a cancer therapy.

## ACKNOWLEDGEMENTS/FUNDING

We thank all members of the Jain lab for thoughtful discussions and review of the manuscript. We thank Longhui Qiu and Mark Looney for catheterizing mice for the *in vivo* tracer experiment. We acknowledge the UCSF Radiopharmaceutical Facility for producing ^18^F-FDG. We acknowledge Benjamin W. Blonder for generating the elevation-by-county code and dataset. We acknowledge Hanifa Mohammed and Annabel Menendez for assistance with generating and validating knockout lines. We thank Brian Plosky for his thoughtful insights during the writing process and Chiara Ricci-Tam and Tami Tolpa for their assistance with graphic design. Sequencing was performed at the UCSF CAT, supported by UCSF PBBR, RRP IMIA, and NIH 1S10OD028511-01 grants. Funding was provided by the National Institute of General Medical Sciences Medical Scientist Training Program, grant T32GM141323 (A.D.M. and D.D.Z.), the National Institute of Diabetes, Digestive, and Kidney Diseases under grant 5F30DK139713 (A.D.M.), the National Science Foundation Graduate Research Fellowship program under grant no. 2034836 (B.T.L.C.), the National Heart, Lung, and Blood Institute K08HL169915 (A.H.B.), the American Heart Association Harold Amos Medical Faculty Development Award (A.H.B.), the National Heart, Lung, and Blood Institute F31HL172581 (W.R.F.), National Institute on Aging R01AG065428 (W.R.F.), the NIH grant T32 GM136547 (J.M.S.), Rombauer Fellowship (I.H.J.), and Keck Medical Research Grant (I.H.J.). H.G. and I.H.J. are Core Investigators at Arc Institute, and their research is supported by Arc. H.G. is also an Era of Hope Scholar (W81XWH-2210121).

Any opinions, findings, and conclusions or recommendations expressed in this material are those of the authors and do not necessarily reflect the views of the National Science Foundation or National Institutes of Health.

## Contributions

I.H.J., H.G., A.D.M., B.T.L.C., and B.M.H.C. conceived the project. B.T.L.C., A.D.M., B.M.H.C., and J.M.S. conducted syngeneic and xenograft tumor experiments, and A.G. assisted with processing syngeneic tumors. A.D.M, B.T.L.C., and A.G.H conducted *in vivo* tumor pO_2_ measurements. B.T.L.C. and A.D.M. bred and conducted experiments with KPC mice, and P.R. performed histopathology on KPC pancreas samples. B.M.H.C. and J.M.S. performed sample preparation for single-cell RNA-sequencing, and C.C. and V.S. processed and analyzed the single-cell transcriptomics data. A.D.M., B.T.L.C., R.S., Y.S., and H.V. conducted ^18^F-FDG experiments. B.T.L.C., A.D.M., W.R.F., and D.D.Z. performed, validated, and tested genetic manipulations. A.D.M and B.T.L.C. designed and conducted tumor and tumor interstitial fluid metabolomics experiments with assistance from S.B., A.G.H., and A.H.B. A.D.M. and B.T.L.C. designed and performed stable isotope-labeled tracer experiments. J.M.S., B.T.L.C., and A.D.M. performed *in vivo* serial reimplantation experiments. B.T.L.C. and A.D.M. conducted chemotherapy experiments with input from B.M.H.C. A.H.B., T.A.J., A.D.M., B.M.H.C., and B.T.L.C. conducted immunotherapy experiments. A.D.M, B.T.L.C., and S.B. performed HypoxyStat experiments. A.D.M., B.T.L.C., H.G., and I.H.J. wrote the manuscript, with input from all authors.

## ETHICS DECLARATIONS

Competing interests: I.H.J., H.G., B.T.L.C., A.D.M., and B.M.H.C. have patents related to hypoxia-based therapies.

## METHODS

### Cell culture

Panc02 and E0771 cells were used for syngeneic tumor models. These cell lines were maintained in DMEM (Thermo Fisher Scientific, 11995073) with 10% FBS (Corning/Fisher Scientific) and 1% penicillin-streptomycin (Gibco/Thermo Fisher Scientific).

For the GENEVA pooled cell line xenograft, HOS143B, SW620, HCC1806, A375, PC3, SJSA1, MDA231, HCT15, LN229, H358, HepG2, Panc1, BxPC3, U87-MG, H441,Hep3b, LNCaP, Caki1, 786O, and SH4 cells were maintained in RPMI (Gibco/Thermo Fisher Scientific) with 10% FBS (Corning), 1% penicillin-streptomycin (Gibco), and 1 µg/mL amphotericin B (Gibco).

To establish GENEVA pools, individual cell line cultures were washed with PBS, trypsinized, and counted using Trypan Blue (Thermo Fisher, 15250061) on an automated cell counter (Bio-Rad, 1450102). Cell lines were combined in a sterile tube at numbers inversely proportional to their growth rates. The pooled cell suspension was centrifuged, the supernatant discarded, and the cell pellet resuspended in undiluted Matrigel (Corning, 356255). All pooling procedures were completed within 2 hours to maintain cell viability.

*In vitro* hypoxia experiments were carried out in C-Chambers (BioSpherix) controlled by a ProOx Model C21 (BioSpherix). Nitrogen gas was generated with a nitrogen generator (N2Gen-02CPi-P, South-Tek Systems). In short, the C-Chamber was housed in a standard tissue culture incubator to warm the chamber to 37 °C. The interior of the chamber was humidified and kept at 5% CO_2_ while nitrogen gas was flowed in until the desired %O_2_ was reached.

### Animals

Mice were housed at room temperature and humidity (approximately 20 °C and 50% humidity) with a 12h:12h light:dark cycle. Mice were fed a chow diet (PicoLab 5058). All experiments with animals were approved by the Institutional Animal Care and Use Committee at UCSF/Gladstone Institutes.

For subcutaneous and orthotopic tumors, either C57BL/6J (The Jackson Laboratory, 000664) or NSG (NOD.Cg-Prkdcscid Il2rgtm1Wjl/SzJ; The Jackson Laboratory, 005557) were used. Mice were anesthetized with 3% isoflurane mixed with oxygen. For syngeneic subcutaneous PDAC tumors, 1 × 10^6^ Panc02 cells in 100 µL of 1:1 matrigel:PBS were injected into the flank of C57BL/6J mice. For syngeneic orthotopic breast cancer tumors, 1 × 10^5^ E0771 cells in 50 µL of 1:1 matrigel:PBS were surgically implanted into the mammary fat pads of C57BL/6J mice. For the anti-CTLA4 antibody experiment, 1 × 10^6^ E0771 cells in 50 µL of 1:1 matrigel:PBS were surgically implanted into the mammary fat pads of C57BL/6J mice.

For the GENEVA pooled cell line xenograft, immunodeficient NSG mice were used (The Jackson Laboratory, 005557). Cell suspensions were prepared at a concentration of 5 × 10^6^ cells per mL and were maintained on ice throughout the injection procedure. Subcutaneous injections were performed bilaterally, with 200 µL of cell suspension administered into each flank.

Tumors were measured using calipers and tumor volumes were estimated using the following formula, where *W* is tumor width and *L* is tumor length: π/6(*W*^2^)(*L*).

For the genetically engineered mouse model of pancreatic cancer, tamoxifen-inducible *Kras^G12D^*;*p53^fl/fl^*;*Pdx1-CreER* mice (The Jackson Laboratory, 032429) were used. Mice that were heterozygous for Kras^LSL-G12D^, homozygous for p53^LoxP^, and homozygous for Pdx1-CreER were bred. Pups were weaned within 3-4 weeks of birth and then dosed with 4 mg of tamoxifen, free base (200 µL at 20 mg/mL, MP Biochemical) for 5 days. 12 weeks after the first dose of tamoxifen, mice were moved to either an 8% O_2_ chamber or maintained in 21% O_2_ as controls. At 16 weeks after the first dose of tamoxifen (4 weeks in hypoxia), mice were sacrificed, and their pancreases were collected and weighed. The mouse and pancreas were kept in 10% neutral buffered formalin (Sigma-Aldrich) for 1 week before being transferred to a 70% ethanol solution. The carcasses and pancreas tissue were then sent for histopathological analysis.

### *In vivo* hypoxia treatment

Hypoxic conditions were created in 472 L acrylic chambers by mixing generated nitrogen (N2Gen-02CPi-P, South-Tek Systems) and compressed air to either 8% or 11% using a gas mixer. The chamber was humidified with open water containers.

The *in vivo* stable isotope labeled tracer experiments were carried out in a glove box (Coy Laboratories). Nitrogen was supplied via a nitrogen generator (N2Gen-02CPi-P, South-Tek Systems) and mixed with oxygen gas (Airgas).

For both setups, the levels of inspired oxygen and carbon dioxide were monitored continuously via a wireless system and verified daily. To prevent CO₂ buildup, soda lime (Fisher Scientific) was placed in each chamber. Animals were kept in their home cage, changed minimum weekly, and were provided water and chow ad libitum.

### Tissue oxygen measurements

Mice were anesthetized with 4% isoflurane and intubated using an intubation platform (Kent Scientific). Once intubated, they were positioned on a heated surface and ventilated with a MiniVent Type 845 (Hugo Sachs, Harvard Apparatus) at a stroke volume of 150 µL and a rate of 150 breaths per minute. Anesthesia was maintained with 2% isoflurane during ventilation.

To control inhaled oxygen levels, pure nitrogen (>99.995%, Airgas) and oxygen (>99.5%, Airgas) were blended using a dual flow meter (Billups-Rothenberg, DFM-3002). Gas cylinders were connected to CGA-580 (nitrogen) and CGA-540 (oxygen) regulators, each set to 5 psig. The oxygen concentration of the gas mixture was monitored with a Flow Through Oxygen Monitor (Billups-Rothenberg, GMS-5002), keeping the inhaled %O₂ within ± 0.2% of the target by adjusting the oxygen flow while maintaining nitrogen flow at 5 L/min. The mixed gas was then routed through an isoflurane vaporizer.

For measuring tissue oxygen levels, a 50 μm Clark-type oxygen electrode (Unisense OX-50) was inserted into tumors following surgical incision. Readings were taken once stabilized for 10 seconds. The electrode was polarized for a minimum of 2 hours before use and calibrated using two standards: room air–saturated PBS at 31°C (159.6 mmHg O₂) and a 31°C anoxic sodium ascorbate solution at pH 9.9 (0 mmHg O₂).

### GENEVA genotyping

Individual cell lines were cultured in RPMI medium (Gibco #11875093) and harvested by washing with PBS followed by trypsinization. Cell suspensions were counted and 1 × 10^6^ cells per line were pelleted by centrifugation. After removing the supernatant, cell pellets were processed for RNA extraction. Total RNA was extracted from the pellets using the Zymo Quick-RNA Microprep Kit (Zymo Research #R1051) according to the manufacturer’s protocol. RNA concentration was assessed using the Qubit HS RNA Assay Kit (Thermo Fisher #Q32852).

For library preparation, 10 ng of total RNA from each cell line was used as input for QuantSeq-Pool 3’ mRNA-Seq Library Prep (Lexogen #139), processed according to manufacturer’s protocol, and sequenced on an Illumina NovaSeq X at a depth of ∼1 × 10^7^ reads per cell line.

The genotyping reads were demultiplexed, trimmed, and UMI-extracted according to the standard QuantSeq-Pool pipeline^77^. Next, they were aligned with STAR^78^ to the same reference used for the single-cell experiment and the unique molecular identifiers (UMIs) were deduplicated using UMI-tools^79^. Reads from each sample were downsampled to the number of reads in the smallest library, so that all samples had the same number of reads. The genotypes were called using BCFtools (1.17)^80^.

### GENEVA pooled tumor collection and dissociation

Tumors were surgically excised and mechanically dissociated using a razor blade. Each tumor was enzymatically digested in 5 mL DMEM (Thermo Fisher, 119650921) containing 0.5 vial Liberase TL (Millipore, 05401020001), 0.5 vial DNase I (Worthington, LK003170), and 32 mg Type 3 Collagenase (Worthington, LS004180) for 45 minutes at 37 °C with continuous agitation.

Following digestion, cells were treated with ACK lysis buffer (Thermo Fisher, A1049201) for 10 minutes at room temperature to lyse red blood cells. After centrifugation, cells were resuspended in PBS and filtered through a 70 μm cell strainer. Single-cell suspensions were counted using Trypan Blue (Thermo Fisher, 15250061) on an automated cell counter (Bio-Rad, 1450102).

2 tumors per oxygen condition were selected for single-cell RNA-sequencing, based on cell yield and viability. 400 HEK cells were spiked into each sample for normalization purposes, and the samples were loaded onto individual lanes of a Chromium NEXT GEM 3’ HT chip (10x Genomics #PN-1000372) at 40,000 cells per lane.

### GENEVA data analysis

We generated GENEVA libraries from the GEMs following the standard Chromium NEXT GEM 3’ HT workflow (10x Genomics, PN-1000372). Based on initial cell loading numbers, we targeted a sequencing depth of 25,000 reads per cell. Sequencing was performed at the UCSF Center for Advanced Technology on an Illumina NovaSeq X.

The single-cell reads were aligned and quantified with cellranger (version 7.1.0)^81^, first to a combined human-mouse reference, and then to a human only reference (refdata-gex-GRCh38-2020-A/GENCODE v32). The cells that aligned to >80% human were then assigned as human cells, and the data from these cells aligned to a human reference was used for all further analysis.

The single cell libraries were demultiplexed with demuxlet^82^. The cells marked as ‘ambiguous’ by demuxlet were assigned a genotype using a custom algorithm that assigned them to the same genotype as the most common genotype as their 7 nearest neighbors. Cells were then filtered using scanpy (version 1.9.5)^83^ for those with >4000 genes and <10% UMIs from mitochondrial genes.

To generate survival data in hypoxia by cell line, the counts of each cell line in hypoxia and normoxia were calculated and averaged by replicate. The proportion of each cell line in the final tumor was divided by the proportion of each cell line in the initial pool of cells that was implanted into mice. This value was multiplied by the average tumor volume in each condition to estimate tumor growth for each cell line. Relative fitness in normoxia over hypoxia was calculated for each cell line as the Log_2_(tumor growth estimate in normoxia/tumor growth estimate in hypoxia) (**Supp. Table 2**).

For each cell line, differentially expressed genes were calculated between hypoxia and normoxia using DESeq2^84^ using the standard Wald test, without fold change shrinkage. Cell cycle counts for each condition were calculated using the scanpy ‘score genes cell cycle’ function, using the cell cycle genes defined in Tirosh et. al^85^. Pearson correlations were calculated between differential expression of each gene and relative fitness in hypoxia. Gene set enrichment scores were calculated on differentially expressed genes using the implementation of GSEA in decoupleR^86,87^. Enrichment analysis among ranked correlation coefficients was conducted using iPAGE^88^.

### Histopathology

To evaluate the presence of neoplastic cells, five serial sections of the whole pancreas and representative longitudinal sections of liver were embedded in paraffin and serially sectioned at 4 μm for staining with hematoxylin and eosin (H&E). Histological analysis was performed by a board-certified comparative pathologist blinded to experimental groups (P.R.). H&E images were analyzed using Olympus cellSens software. Tumors were graded according to previously published criteria^89^. Briefly, tumors were classified as pancreatic intraductal neoplasia (PanIN) or invasive carcinoma based on cell morphology, tissue architecture and organ invasion. Staging of the tumors was performed using the “tumor, node, metastasis” (TNM) staging scheme^90^. The liver was the sentinel organ used to evaluate the presence of metastasis.

### Blood glucose measurements

Non-fasted blood glucose measurements were conducted from tail blood samples using a portable glucometer (AimStrip Plus).

### ^18^F-FDG preparation

[¹⁸F]fluoride ion was generated via the ¹⁸O(p,n)¹⁸F nuclear reaction by bombarding oxygen-18 enriched water (Rotem) with protons using a GE PETtrace™ cyclotron. ^18^F-FDG was synthesized in the UCSF Radiopharmaceutical Facility using the [¹⁸F]fluoride ion and an automated GE FASTlab system equipped with the FDG Citrate FASTlab reagent cassette (GE).

### ^18^F-FDG injection & gamma radiation quantification

FDG injections and gamma radiation quantification were conducted at the Preclinical Imaging Core facility at China Basin in the UCSF Department of Radiology & Biomedical Imaging. During transport, mice were housed in enclosed boxes, and portable nitrogen and oxygen tanks (Airgas) were used to maintain the desired FiO₂ levels. Anesthesia was induced with 3% isoflurane, and FDG was administered via tail vein. After a 60-minute uptake period, mice were euthanized, and tumors were dissected and weighed.

Gamma radiation levels in each tumor were quantified using a HIDEX Automatic Gamma Counter (HIDEX). Radioactivity levels were normalized to both organ weight and the administered dose at the time of counting. The percent injected dose per gram (%ID/g) was calculated. The remaining radioactive dose at the time of quantification was decay-corrected based on the known half-life of the radiotracer, using the following equation, where *Δt* denotes the time elapsed in minutes between injection and counting:

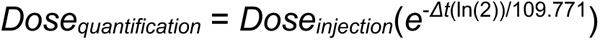

### Genetic deletions

The lentiCRISPRv2 backbone plasmid (AddGene 52961) was digested and ligated with sgRNAs. The resulting plasmid was packaged into lentivirus by transfecting the plasmid into HEK293T/17 cells along with the packaging plasmids pVSVg (AddGene 8454) and psPAX2 (AddGene 12260). The resulting lentiviral particles were transduced into Panc02 cells by spinfection, and cells were selected in culture media containing 4 µg/mL puromycin for 2 days followed by 8 µg/mL puromycin for 2 days. Single clones were plated in 96-well plates and expanded until knockouts could be validated.

### Immunoblotting

Cells were lysed with ice-cold RIPA buffer (Thermo Fisher Scientific) supplemented with Halt Protease and Phosphatase Inhibitor Cocktail. Protein abundance was quantified using a Pierce BCA Protein Assay Kit (Thermo Fisher Scientific) and equalized across samples. 6X Laemmli SDS sample buffer (Thermo Fisher Scientific) was added to each lysate, and samples were boiled at 95 °C for 5 minutes. Protein lysates were run on SDS-PAGE gels (Bio Rad Mini-PROTEAN TGX) for 45 minutes at 150 V. Proteins were transferred to PVDF membranes using Trans-Blot Turbo (Bio Rad), and membranes were blocked using 5% non-fat milk in TBST (Thermo Fisher Scientific) for 1 hour. Primary antibodies were diluted 1:1000 in 3% non-fat milk in TBST, and membranes were incubated with primary antibody solutions overnight at 4 °C. After washing with TBST, membranes were incubated with a 1:5000 solution of secondary antibodies in 5% non-fat milk in TBST for 1 hour at room temperature. Bands were visualized using ECL (Thermo Fisher Scientific).

### RT-qPCR

Panc02 cells in 24-well plates were lysed in 350 µL of RLT buffer with 1% β-mercaptoethanol. RNA was extracted using the RNeasy Mini Kit (Qiagen) and reverse transcribed using the QuantiTect Reverse Transcription Kit (Qiagen). RT-qPCRs were conducted by combining the cDNA with the Maxima SYBR Green/ROX qPCR Master Mix (Thermo Fisher Scientific) and primers for each gene of interest. Relative expression was determined using the ΔΔCT method, and expression of each gene was normalized to *β-actin*.

### Tumor polar metabolite extraction

Upon dissection, tumors were flash frozen in liquid nitrogen and stored at −80 °C. Frozen tumors were pulverized using a Cryo-Cup grinder (BioSpec Products), and the resulting powder was weighed. For each mg of tumor homogenate, 20 µL of 40/40/20 methanol/acetonitrile/water with 1 mM D8-valine as an internal standard was added. Samples were mixed in a thermomixer at 4 °C, sonicated for 3 minutes, and immersed in liquid nitrogen, and this process was repeated. Samples were then incubated on ice for 20 minutes and centrifuged at 20,000 g for 20 minutes at 4 °C. Supernatants were collected and lyophilized overnight. The lyophilized tumor extracts were resuspended in 60% acetonitrile, and the samples were sonicated for 3 minutes, mixed in a thermomixer at 4 °C, and incubated for 20 minutes on ice. Samples were centrifuged at 20,000 g for 20 minutes at 4 °C, and the supernatant was transferred to autosampler vials for analysis.

### Tumor interstitial fluid polar metabolite extraction

Tumor interstitial fluid was isolated from subcutaneous tumors as previously described ^65,91^. Nylon mesh filters were affixed over 50 mL conical tubes, allowing sufficient flexibility for tumor compression. Prior to dissection, 25 mL of pre-chilled (4 °C) PBS was aliquoted into petri dishes for tumor washing, and >4 cm square pieces of Whatman paper were prepared for blotting. Mice were euthanized, and tumors were excised. Each tumor was washed in PBS, thoroughly blotted dry using fresh Whatman paper to remove residual buffer, and placed on the prepared nylon mesh atop conical tubes. Tumors were either processed immediately or stored briefly on ice. Tumors were centrifuged at 4 °C for 10 minutes at 100 g to collect tumor interstitial fluid (TIF). Some samples required additional centrifugation: either an additional 5 minutes at 100 g or, in the case of certain smaller tumors, an additional 5 minutes at 200 g. TIF samples were collected into pre-labeled tubes and stored on dry ice. Remaining tumors were transferred into labeled tubes and flash-frozen in liquid nitrogen. All TIF samples were maintained on dry ice until storage.

Polar metabolites were extracted by adding 20 µL of 90% acetonitrile with 1 mM D8-valine as an internal standard for every 5 µL of TIF. Samples were vortexed and mixed on a thermomixer for 5 minutes at 4 °C, and then they were incubated on ice for 20 minutes. Finally, they were centrifuged at 20,000 g for 20 minutes at 4°C, and the final supernatants were transferred to autosampler vials.

### Plasma polar metabolite extraction

Plasma was isolated by centrifuging whole blood at 2,000 g for 10 minutes at 4 °C and collecting the resulting supernatant. Polar metabolites were extracted by adding 45 μL of 80% methanol to 5 μL of plasma. The mixture was vortexed for 10 seconds and then incubated at −80 °C for 20 minutes. Samples were subsequently centrifuged at 20,000 g for 10 minutes at 4 °C, and the supernatants were collected. These were lyophilized at 4 °C and reconstituted in 40 μL of 60% acetonitrile. Reconstituted samples were sonicated, mixed in a thermomixer, and incubated at 4 °C for 20 minutes. After a final centrifugation at 20,000 g for 20 minutes at 4 °C, the clarified supernatants were transferred to appropriate vials for liquid chromatography-mass spectrometry (LC-MS) analysis.

### Stable isotope labeled tracer experiments in cell culture

Stable isotope-labeled DMEM was prepared using a base DMEM without glucose, glutamine, isoleucine, leucine, valine, sodium pyruvate, sodium bicarbonate, and phenol red (US Biologicals, D9800-36). Each missing component was added to the base DMEM but ɣ-^15^N-glutamine (Cambridge Isotope Laboratories, NLM-557) was used instead of regular glutamine. 10% dialyzed FBS and 1% penicillin-streptomycin were added, and the final media was supplemented with 10 µM 8-^13^C-adenine (MedChemExpress, HY-B0152S1).

6 × 10^5^ Panc02 cells were plated in 6-well plates and incubated at either 21% O_2_ or 0.5% O_2_. After 18 hours of incubation, cells were washed with PBS and cultured in complete DMEM (including glucose and pyruvate) containing 4 mM ɣ-^15^N-glutamine and were supplemented with 10 µM of 8-^13^C-adenine. Six hours post-treatment, culture media was removed and cells were washed with PBS. Cells were then quenched with 1 mL of ice-cold 80% methanol. Plates were incubated at −80 °C for 30 minutes. Cells were scraped and transferred into Eppendorf tubes, vortexed briefly, and incubated on ice for 20 minutes. Samples were centrifuged at 20,000 g for 20 minutes at 4 °C. The supernatant was transferred to fresh tubes and lyophilized overnight. Lyophilized samples were resuspended in 60% acetonitrile. Samples were sonicated for 3 minutes, mixed in a thermomixer at 4 °C, and incubated for 20 minutes on ice. Samples were centrifuged at 20,000 g for 20 minutes at 4 °C, and the supernatant was transferred to autosampler vials for analysis.

### Stable isotope labeled tracer experiments in mice

Catheters (Instech, C20PU-MJV2014) were surgically placed in the right jugular veins of mice and were connected to a vascular access button (Instech, VABM1B/25) on the backs of the mice. An infusion solution containing 283 mM ɣ-^15^N-glutamine and 4 mM 8-^13^C-adenine in 0.9% saline (Fisher Scientific, Z1376) was prepared^44^. Mice were connected to an infusion line on a counter-balanced lever arm (Instech, SMCLA) to allow free movement, and the infusion line was connected to a syringe pump (Harvard Apparatus, 70-4501). Following a 150 µL/minute priming dose for 1.5 minutes, the infusion rate was lowered to 2.5 µL/minute for 5 hours. At the end of the infusion, blood was collected by cardiac puncture, mice were euthanized, and tumors were dissected and flash frozen.

### Liquid chromatography-mass spectrometry

Samples were analyzed using a high-resolution Orbitrap Exploris 240 mass spectrometer (Thermo Fisher Scientific) equipped with electrospray ionization. The mass spectrometer was coupled to a Vanquish Horizon ultra-high performance liquid chromatography system using hydrophilic interaction chromatography. A 2 μL aliquot of each polar metabolite extract was injected and separated on an iHILIC-(P) Classic column (2.1 × 150 mm, 5 µm; HILICON AB, 160.152.0520). The autosampler was maintained at 4 °C, and the column temperature was set to 40 °C. The mobile phase consisted of 20 mM ammonium bicarbonate in water (adjusted to pH 9.6 with ammonium hydroxide) as phase A, and acetonitrile as phase B. Chromatographic separation was performed at a flow rate of 200 μL/min with the following gradient: 0–20 min, 80% to 20% B; 20–20.5 min, linear ramp from 20% to 80% B; 20.5–28 min, held at 80% B. A 10-minute equilibration at initial conditions preceded each run.

The mass spectrometer operated in full-scan, polarity-switching mode with a spray voltage of 3.5 kV in positive mode and 3.25 kV in negative mode. Spectra were acquired from m/z 70 to 1000 at a resolution of 60,000. Additional settings included an RF Lens level of 60%, AGC target of 1e7, and a maximum injection time of 200 ms. The sheath gas was set to 35 units, auxiliary gas to 10 units, and sweep gas to 0.5 units. The ion transfer tube and vaporizer temperatures were 300 °C and 35 °C, respectively.

### Semi-targeted metabolomics

For semi-targeted metabolomics, the Mass Spectrometry Metabolite Library of Standards (MSMLS, IROA Technologies) was analyzed on the LC-MS system. Fragmentation spectra were collected by running the standards in both positive and negative ion modes using data-dependent MS2 acquisition. HCD collision energies of 15%, 45%, and 90% were applied, and spectra were acquired at a resolution of 30,000 on the Orbitrap. For compound identification, pooled quality control samples were prepared by combining equal volumes from each experimental sample. QC samples were analyzed using the same ddMS2 settings and a targeted mass list derived from detectable MSMLS compounds. Additionally, untargeted compound identification was performed using AcquireX with identical ddMS2 parameters.

Compound identification and peak integration were carried out using Compound Discoverer (Thermo Fisher Scientific), referencing both a custom spectral library built from MSMLS standards and the online mzCloud database.

### Serial reimplantation of Panc02 tumors

Each mouse received bilateral subcutaneous flank injections of 100 µL containing 1 × 10^6^ cells per injection site. After approximately one month of growth, tumors were harvested, dissociated into single-cell suspensions, and cultured in vitro for 4-7 days before reimplantation into naive mice. This serial transplantation process was repeated for four passages.

### *In vivo* drug treatments

Gemcitabine (Millipore Sigma, 1288463) was dissolved in 0.9% saline (Fisher Scientific, Z1376) and dosed at 100 mg/kg via IP injection. For this experiment, mice in the hypoxia arm were placed in 11% O_2_ for 1 week prior to tumor injection to allow them to acclimate to hypoxia as stated in the IACUC protocol. Subcutaneous Panc02 tumors were injected as described above. Mice in the hypoxia arm were placed into 8% O_2_. Once tumors were >100 mm^3^, mice were either dosed with gemcitabine or saline. After 32 days, the frequency of gemcitabine dosing was increased to twice per week.

Anti-Ctla4 and isotype IgG-2b antibodies (BioXCell) were diluted to 1 µg/µL in dilution buffer (BioXCell). Upon tumor implantation, mice were either moved to 8% O_2_ or kept in 21% O_2_ and dosed with 100 µg anti-Ctla4 or isotype control antibody 3, 6, 9, and 12 days after implantation.

Mice were dosed with either 400 mg/kg or 600 mg/kg HypoxyStat. HypoxyStat (Wuxi AppTech) was formulated as a suspension at 40 mg/mL or 60 mg/mL in 0.5% hydroxypropyl methylcellulose (Sigma) and stored at 4 °C with agitation. 24 hours after tumor implantation, mice were dosed via oral gavage with 400 mg/kg HypoxyStat or vehicle daily for 1 week and 600 mg/kg HypoxyStat or vehicle daily thereafter. Mice in the 8% arm were dosed daily with vehicle. All mice were weighed daily to determine the dose of HypoxyStat or vehicle to administer.

### Cancer epidemiology

Mean county elevation was calculated by running zonal statistics on the 2018 Cartographic Boundary Shapefiles (United States Census Bureau) and the 2022 ETOPO Global Relief Model (60-Arc second resolution Geoid height, National Centers for Environmental Information, National Oceanic and Atmospheric Administration) in R (4.3.2)^92^ using the following packages: sf (1.0-21)^93,94^, dplyr (1.1.4)^95^, and terra (1.8-60)^96^.

The per county, age-adjusted cancer mortality rate used data from the U.S. Cancer Statistics database, which integrates information from two primary sources: the National Program of Cancer Registries (NPCR) of the Centers for Disease Control and Prevention (CDC) and the Surveillance, Epidemiology, and End Results (SEER) Program of the National Cancer Institute. The dataset encompasses cancer incidence information from NPCR-affiliated central cancer registries in 46 states and the District of Columbia, along with SEER registry data from four additional states^97^. Average annual cancer rates for the period 1999–2020 were calculated per 100,000 people and age-adjusted to the 2000 U.S. standard population using the direct method with 19 age groups^98^. Rates derived from fewer than 16 cases were excluded.

The county level obesity and diabetes data used in the regression was from the CDC’s U.S. Diabetes Surveillance System 2018 data. The county level median household income data used in the regression is from the U.S. Census Bureau’s American Community Survey and is the 5-year data for 2021. The median income values are in 2021 inflation adjusted dollars. All epidemiological, zonal statistic, and regression analyses were performed in R (4.3.2)^92^. In addition to the packages listed above, the following were used: ggplot2 (3.5.2)^99^, readxl (1.4.5)^100^, usmap (0.8.0)^101^, readr (2.1.5)^102^, stringr (1.5.1)^103^, and usmapdata (0.6.0)^104^.

