## Supplemental Figures for "Systemic hypoxia suppresses solid tumor growth"

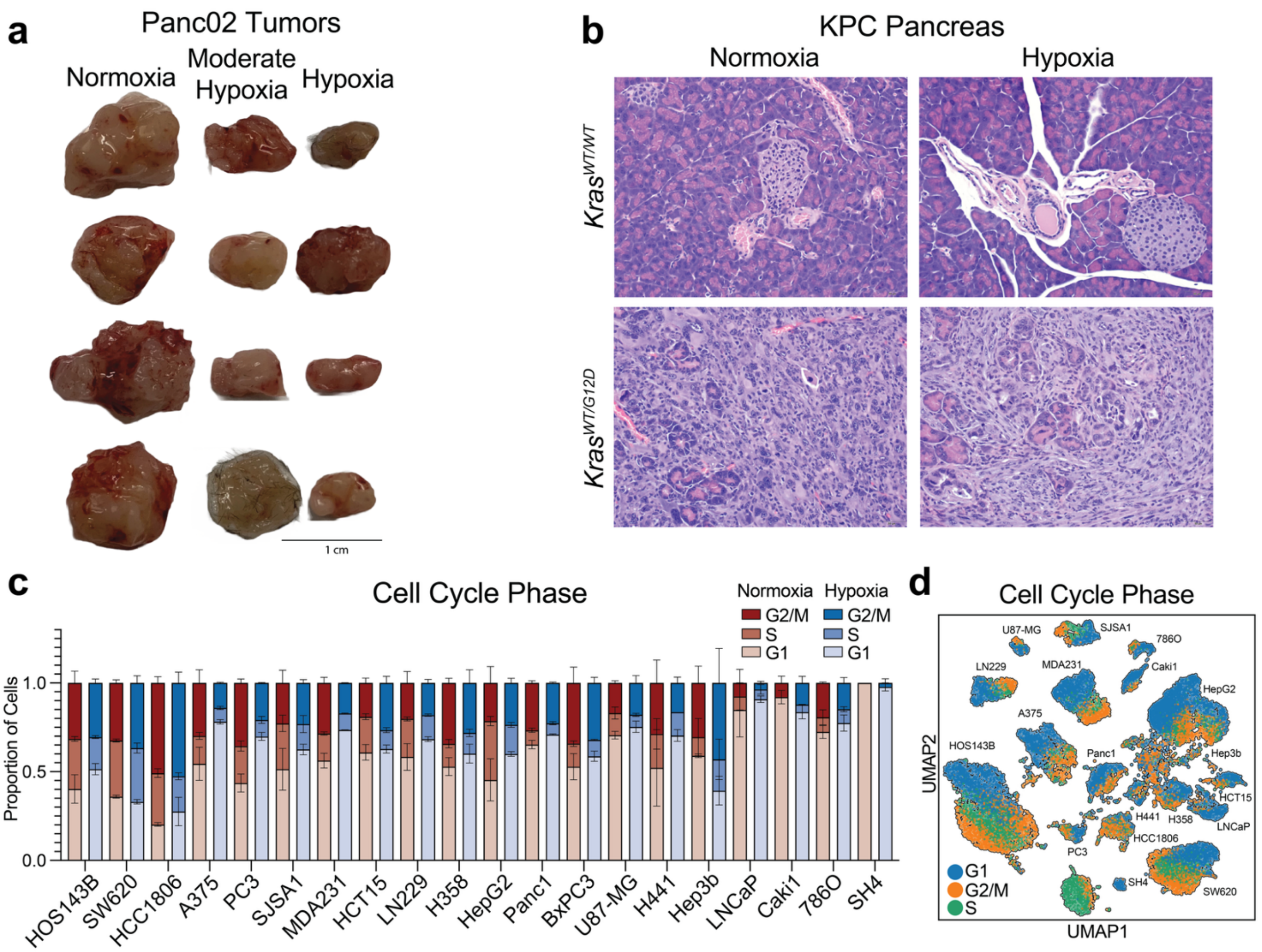


**Figure S1. Hypoxia suppresses primary tumor growth in mouse PDAC. a,** Representative cropped images of tumors dissected from mice housed in normoxia (21% $O_2$), moderate hypoxia (11% $O_2$), or hypoxia (8% $O_2$). **b,** Representative histopathology images of pancreases from KPC mice, either wild-type *Kras* or heterozygous for the *Kras*$^{G12D}$ mutation, and exposed to normoxia or hypoxia. **c,** Proportion of each cell line from the GENEVA experiment in G1, S, or G2/M phase from mice exposed to normoxia or hypoxia, determined based on single-cell transcriptomics. $n$ = 2 tumors per group. **d,** UMAP projection of single-cell transcriptomics from the GENEVA experiment with cells colored by cell cycle phase.

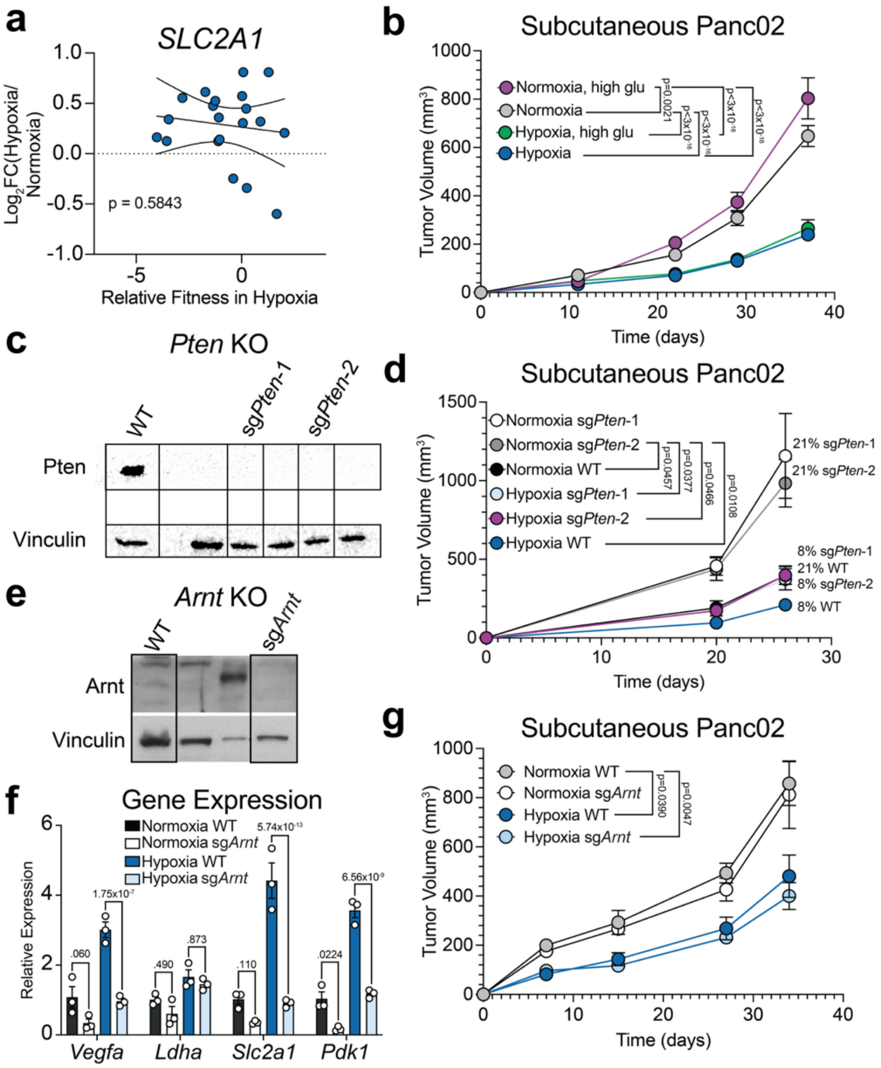


**Figure S2. Tumor suppression in hypoxia is independent of glucose, insulin signaling, and HIF activation. a,** Correlation between fold change in *SLC2A1* expression between hypoxia (8% $O_2$) and normoxia (21% $O_2$) and the relative fitness in hypoxia of each cell line from GENEVA tumors. **b,** Growth of Panc02 subcutaneous tumors in mice that were housed in normoxia or hypoxia and given either normal water or water with 30% glucose. Hypoxia with 30% glucose $n$ = 16 mice, every other group $n$ = 14 mice. **c,** Western blot of Pten protein levels in wild-type cells and in sg*Pten* clones. **d,** Growth of wild-type or sg*Pten* Panc02 subcutaneous tumors in mice that were housed in normoxia or hypoxia. Normoxia sg*Pten*-1, hypoxia sg*Pten*-2 $n$ = 7 mice, every other group $n$ = 8 mice. **e,** Western blot of Hif-1β protein levels in wild-type cells and sg*Arnt* clones. **f,** Relative expression of Hif target genes in wild-type and sg*Arnt* clones after 24 hours of incubation in normoxia (21% $O_2$) or hypoxia (1.5% $O_2$). $n$ = 3 replicates. **g,** Growth of wild-type or sg*Arnt* Panc02 subcutaneous tumors in mice that were housed in normoxia (21% $O_2$) or hypoxia (8% $O_2$). Data are presented as mean ± SEM. Comparisons were made using two-way ANOVA followed by Tukey's multiple comparisons test.

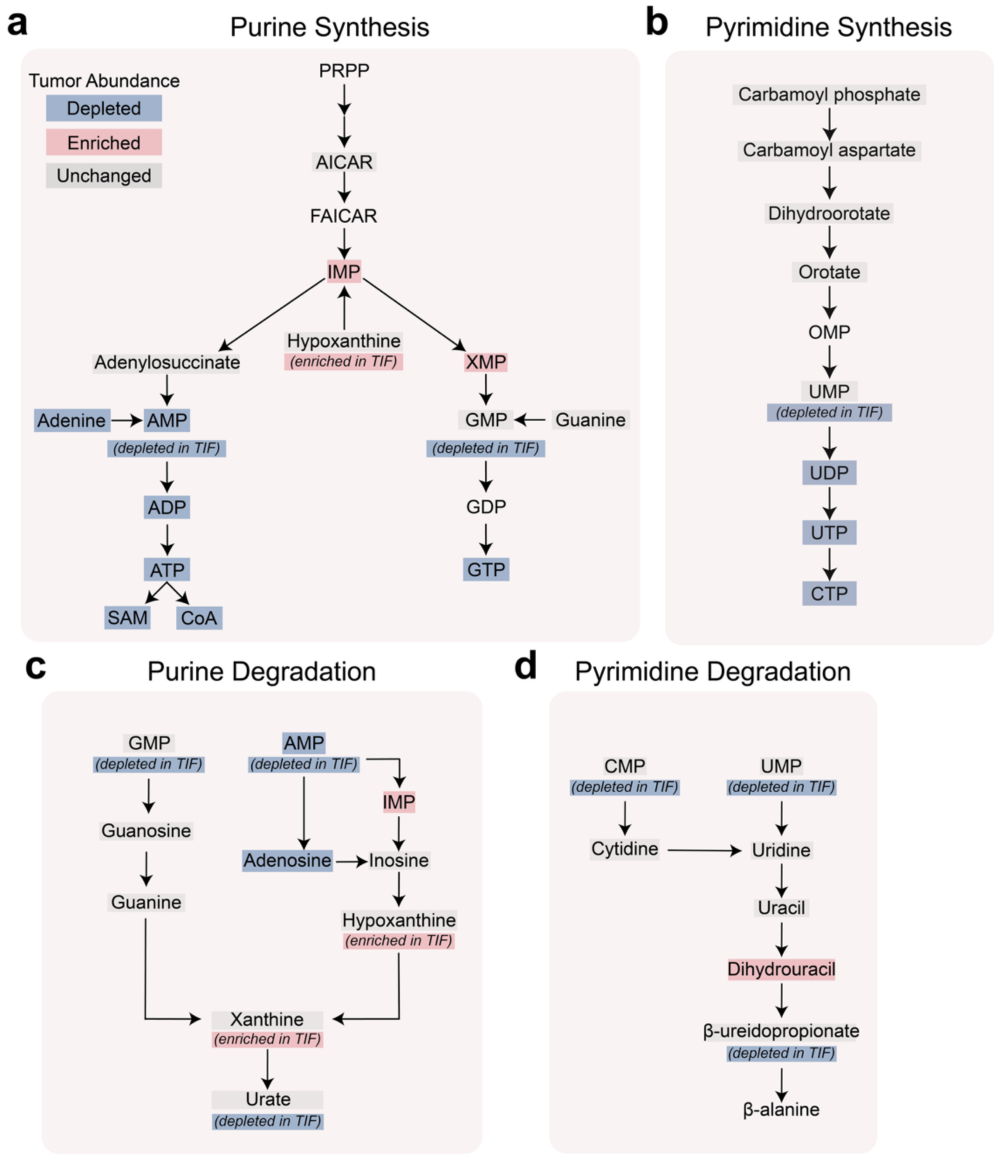


**Figure S3. Hypoxic tumors undergo reprogramming of nucleotide metabolism. a-d,** Schematics of purine synthesis **(a)**, pyrimidine synthesis **(b)**, purine degradation **(c)**, and pyrimidine degradation **(d)** pathways. Metabolites that were depleted in tumors from hypoxic mice (8% $O_2$) compared to normoxic mice (21% $O_2$) are highlighted in blue, and metabolites that were enriched in tumors from hypoxic mice are highlighted in red. Metabolites whose abundance changed between normoxia and hypoxia in TIF are annotated.

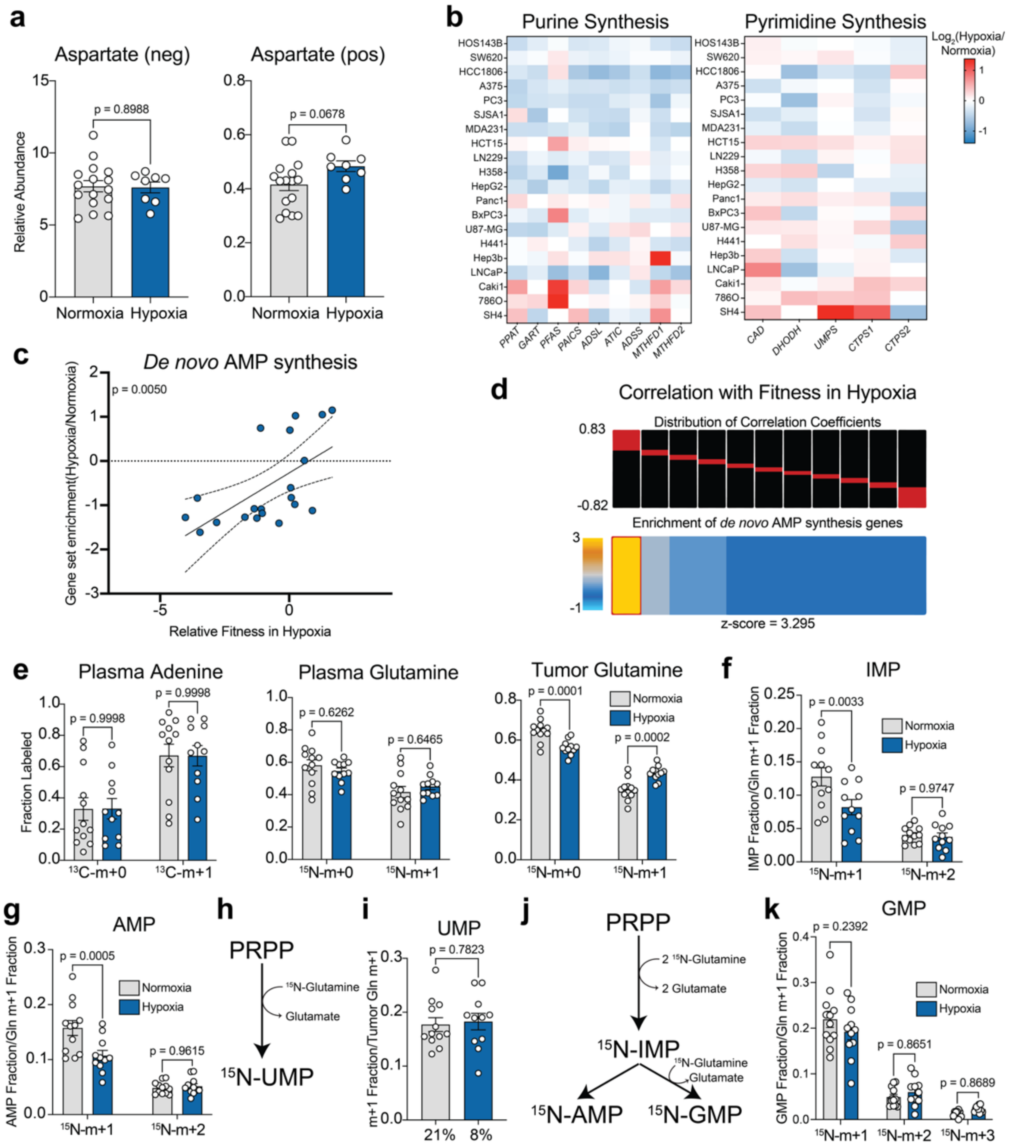


**Figure S4. Systemic hypoxia suppresses de novo purine synthesis**

**a,** Relative abundance (peak area normalized to an internal standard) of aspartate in tumors from mice housed in normoxia (21% $O_2$) or hypoxia (8% $O_2$). Normoxia *n* = 16 mice, hypoxia *n* = 8 mice. **b,** Fold change in expression of *de novo* purine and pyrimidine synthesis genes in different cancer cell lines from GENEVA tumors that were implanted into mice housed in normoxia or hypoxia. **c,** Correlation between the change in GSEA normalized enrichment scores for *de novo* AMP synthesis genes between hypoxia and normoxia and relative fitness in hypoxia. **d,** Enrichment pattern of *de novo* AMP synthesis genes in the distribution of correlation coefficients. Correlation coefficients were calculated for each gene between its change in expression in hypoxia in each cell line and the relative fitness of each cell line in hypoxia. The top row shows binned distributions of correlation coefficients in red, and the bottom row shows over- or under-representation of *de novo* AMP synthesis genes in each bin. **e,** Fractional labeling of plasma $^{13}$C-adenine, plasma $^{15}$N-glutamine, and intra-tumor $^{15}$N-glutamine in mice housed in normoxia or hypoxia. Normoxia *n* = 12 mice, hypoxia *n* = 11 mice. **f-g,** Normalized fractional labeling of each isotopomer (m+1 and m+2) of intra-tumor $^{15}$N-IMP **(f)** and AMP **(g)** normalized to intra-tumor fractional $^{15}$N-glutamine labeling. **h,** Schematic summarizing incorporation of the $^{15}$N label from ɣ-$^{15}$N-glutamine into UMP. **i,** Fractional labeling of intra-tumor $^{15}$N-UMP normalized to intra-tumor $^{15}$N-glutamine fractional labeling in mice housed in normoxia or hypoxia. Normoxia *n* = 12 mice, hypoxia *n* = 11 mice. **j,** Schematic summarizing incorporation of the $^{15}$N label from ɣ-$^{15}$N-glutamine into GMP. **k,** Normalized fractional labeling of each isotopomer (m+1, m+2, and m+3) of intra-tumor $^{15}$N-GMP normalized to intra-tumor fractional $^{15}$N-glutamine labeling. Data are presented as mean ± SEM. Comparisons were made using two-tailed Student's t-tests **(a, i)** or two-way ANOVA followed by Šídák's multiple comparisons test **(e-g, k)**. The z-score for enrichment of *de novo* AMP synthesis genes was calculated using iPAGE **(d)**.

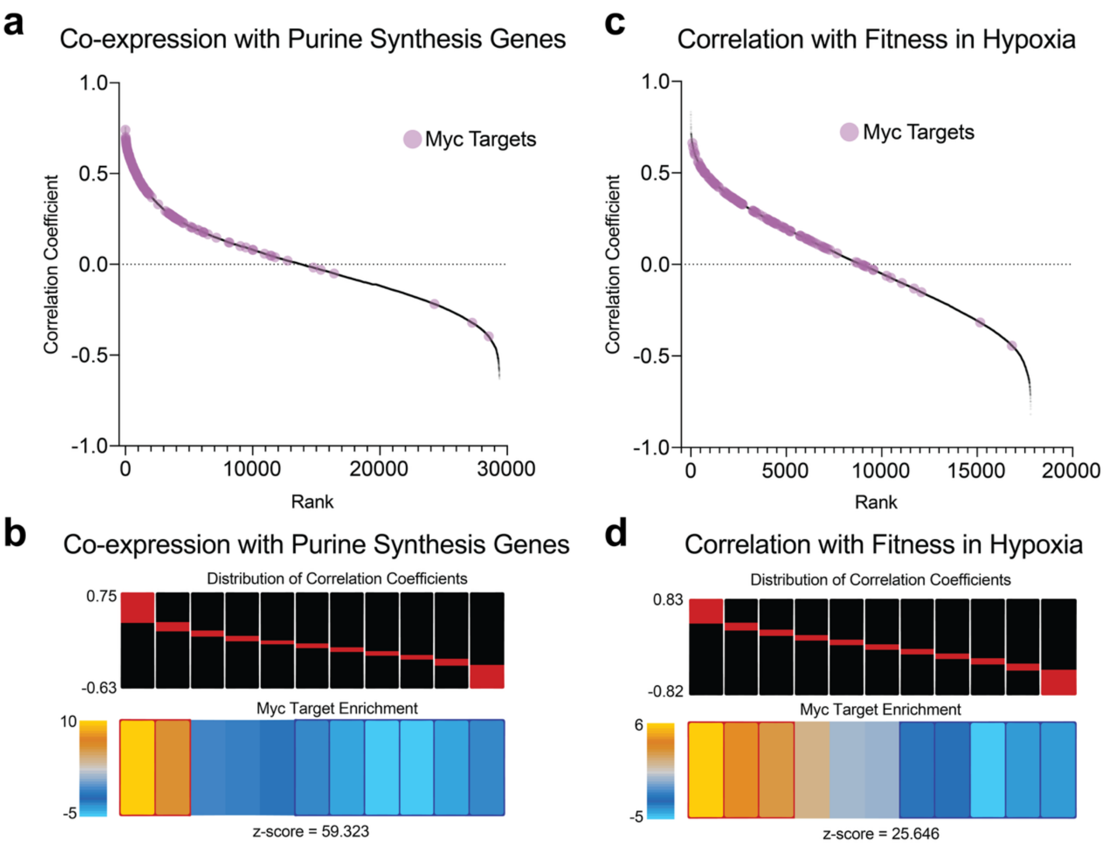


**Figure S5. Myc target expression is associated with expression of *de novo* purine synthesis genes and fitness in hypoxia**
**a,** Ranked correlation coefficients between fold change in a given gene's expression and fold change in expression of the *de novo* purine synthesis genes *PPAT*, *MTHFD1*, *MTHFD2*, *PAICS*, and *ATIC*. Myc targets are labeled in purple. **b,** Enrichment pattern of Myc target genes in the distribution of correlation coefficients shown in **(a)**. The top row shows binned distributions of correlation coefficients in red and the bottom row shows over- or under-representation of Myc targets in each bin. **c,** Ranked correlation coefficients between fold change in a given gene's expression and relative fitness in hypoxia. Myc targets are labeled in purple. Correlation coefficients were calculated using Pearson correlation. **d,** Enrichment pattern of Myc target genes in the distribution of correlation coefficients shown in **(c)**. Z-scores were calculated using iPAGE.

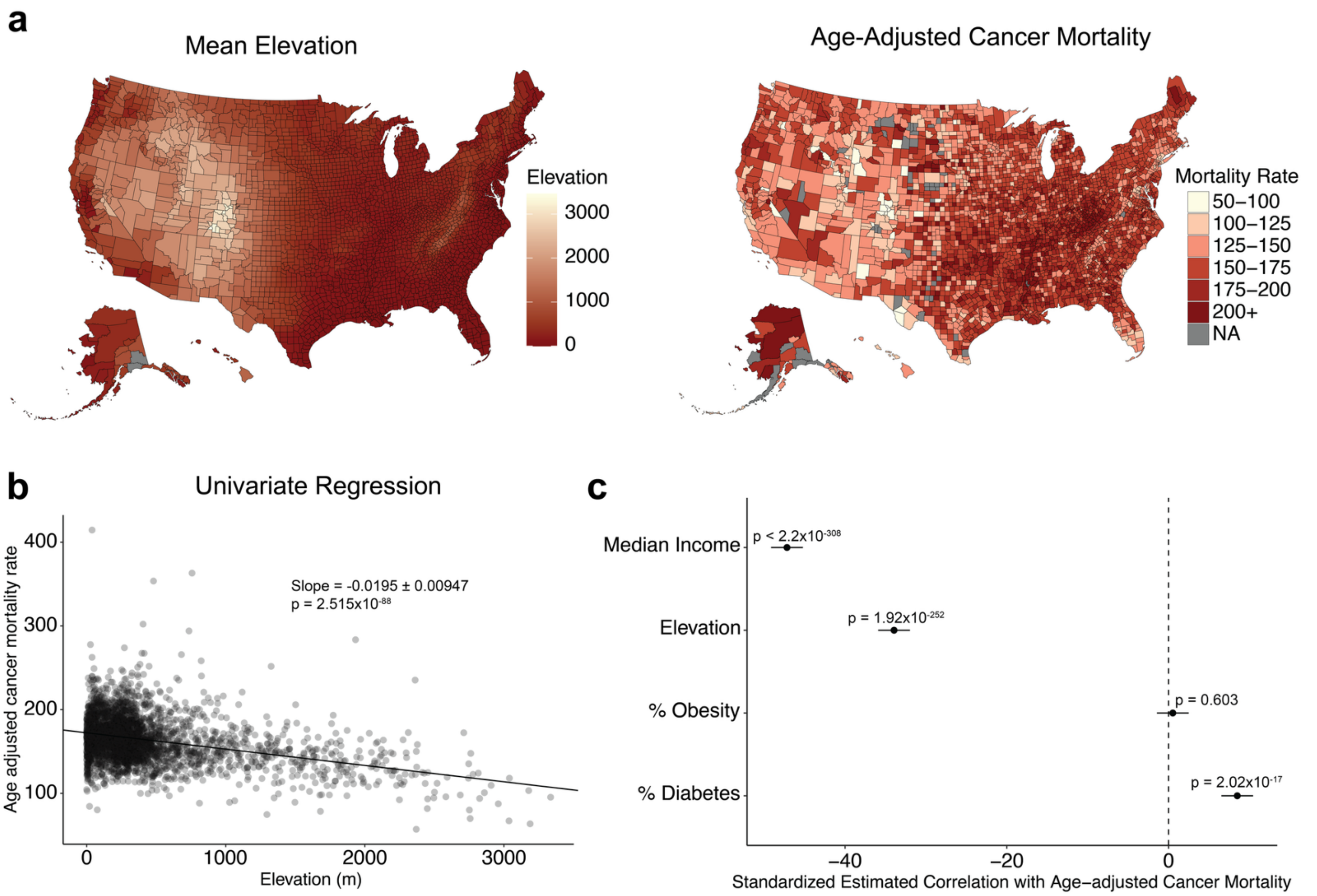


**Figure S6. High-altitude counties across America have lower age-adjusted cancer mortality**

**a,** Heatmaps showing mean elevation in meters and age-adjusted cancer mortality in each county in the USA. **b,** Scatterplot showing the relationship between age-adjusted cancer mortality rate and mean elevation across the USA. **c,** Forest plot showing the standardized estimated correlation coefficients of multiple variables with age-adjusted cancer mortality with a 95% confidence interval. Correlation coefficients were calculated using a generalized linear model.
